# A Temporal and Spatial Atlas of Adaptive Immune Responses in the Lymph Node Following Viral Infection

**DOI:** 10.1101/2025.01.31.635509

**Authors:** Shaowen Jiang, Madhav Mantri, Viviana Maymi, Scott A. Leddon, Peter Schweitzer, Subash Bhandari, Chase Holdener, Ioannis Ntekas, Christopher Vollmers, Andrew I. Flyak, Deborah J. Fowell, Brian D. Rudd, Iwijn De Vlaminck

## Abstract

The spatial organization of adaptive immune cells within lymph nodes is critical for understanding immune responses during infection and disease. Here, we introduce AIR-SPACE, an integrative approach that combines high-resolution spatial transcriptomics with paired, high-fidelity long-read sequencing of T and B cell receptors. This method enables the simultaneous analysis of cellular transcriptomes and adaptive immune receptor (AIR) repertoires within their native spatial context. We applied AIR-SPACE to mouse popliteal lymph nodes at five distinct time points after Vaccinia virus footpad infection and constructed a comprehensive map of the developing adaptive immune response. Our analysis revealed heterogeneous activation niches, characterized by Interferon-gamma (IFN-γ) production, during the early stages of infection. At later stages, we delineated sub-anatomical structures within the germinal center (GC) and observed evidence that antibody-producing plasma cells differentiate and exit the GC through the dark zone. Furthermore, by combining clonotype data with spatial lineage tracing, we demonstrate that B cell clones are shared among multiple GCs within the same lymph node, reinforcing the concept of a dynamic, interconnected network of GCs. Overall, our study demonstrates how AIR-SPACE can be used to gain insight into the spatial dynamics of infection responses within lymphoid organs.

## INTRODUCTION

The adaptive immune system, mediated primarily by T and B cells, is central to pathogen defense, immune regulation, and the establishment of long-term immunological memory^1^. T and B cells recognize specific antigens via their respective adaptive immune receptors (AIRs), the T cell receptor (TCR) and B cell receptor (BCR). The vast diversity of these receptors arises through V(D)J recombination, junctional diversity, and, in the case of B cells, somatic hypermutation^2,3^. Secondary lymphoid organs serve as key sites where naïve T and B cells encounter antigens, become activated, and undergo clonal expansion and selection^4^. Elucidating the spatial organization of these immune cells within lymphoid organs is critical for understanding the cellular interactions and dynamic processes that drive immune responses during infection and disease.

Lymph nodes (LNs) are highly organized structures that facilitate pathogen defense and the orchestration of adaptive immunity^5,6^. Following infection, LNs not only act as sites for activation of adaptive immune cells but also function as dynamic microenvironments where cellular interactions evolve over time and space. Despite their central role in immune responses, several key aspects of LN biology remain poorly understood: How do innate immune cells initiate and coordinate infection responses within the LN microenvironment^7^? What mechanisms and spatial dynamics govern early T cell activation^8^? How do germinal center (GC) B cells exit the GC and differentiate into antibody-secreting plasma cells^9–12^? And at what specific times and locations does class-switch recombination occur in GC B cells^13^? While imaging^14^, computational modeling^9^, and bulk or single-cell sequencing^10,12^ have yielded valuable insights, these questions have yet to be pursued with high-throughput, spatially resolved molecular analysis.

Recent advances in spatial transcriptomics (ST) enable the analysis of spatial cellular and molecular organization in tissues^15–17^. However, most ST methods rely on short-read sequencing, which is insufficient to resolve full-length TCR and BCR sequences. Several approaches have been proposed to overcome this limitation. Spatial VDJ^18^ leverages the Visium platform (100 µm pixel size) to map AIR transcripts in tissues using both long-read and PCR-based short-read sequencing. Slide-TCR-seq^19^ builds off the higher spatial resolution of Slide-seq (10 µm pixel size) to spatially map TCR transcripts and transcriptomes. However, these methods are limited in resolving AIR clonotypes at single-cell and single-nucleotide resolution or detecting BCRs and TCRs concurrently. Additionally, these approaches have been applied at a single time point, offering only a snapshot of immune cells in one organ.

To address these challenges, we developed AIR-SPACE, a methodology that combines high-resolution spatial transcriptomics with error-corrected, paired, full-length immune receptor sequencing. By applying AIR-SPACE to draining popliteal lymph node (PLN) samples collected at five timepoints following footpad injection with recombinant Vaccinia virus (VACV-gB)^20^, we constructed a spatially resolved molecular atlas of LN immune responses throughout the course of infection. The data revealed the emergence of spatial niches associated with early LN responses, spatial and molecular evidence that plasma cells exit GCs via the dark zone–medulla interface, and evidence for B cell clonal recirculation between GCs. Collectively, these results underscore the potential of AIR-SPACE for dissecting complex immunological processes within lymphoid organs.

## RESULTS

### High-resolution spatial mapping of immune repertoires via AIR-SPACE

We designed and implemented AIR-SPACE, a spatial sequencing approach for simultaneous characterization of transcriptomes and adaptive immune receptor repertoires with high spatial and sequence resolution (**Fig. 1**). This assay offers: ***i)*** High spatial resolution (10 µm), ***ii)*** High sequence resolution, allowing recovery of long-read and paired receptor sequences for both B and T cells with low base calling error rate, and ***iii)*** Unbiased spatial transcriptomics of the same tissue. This is achieved by integrating high-resolution spatial transcriptomics with long-read, error-corrected adaptive immune receptor sequencing (**Methods**).

**Figure 1.**
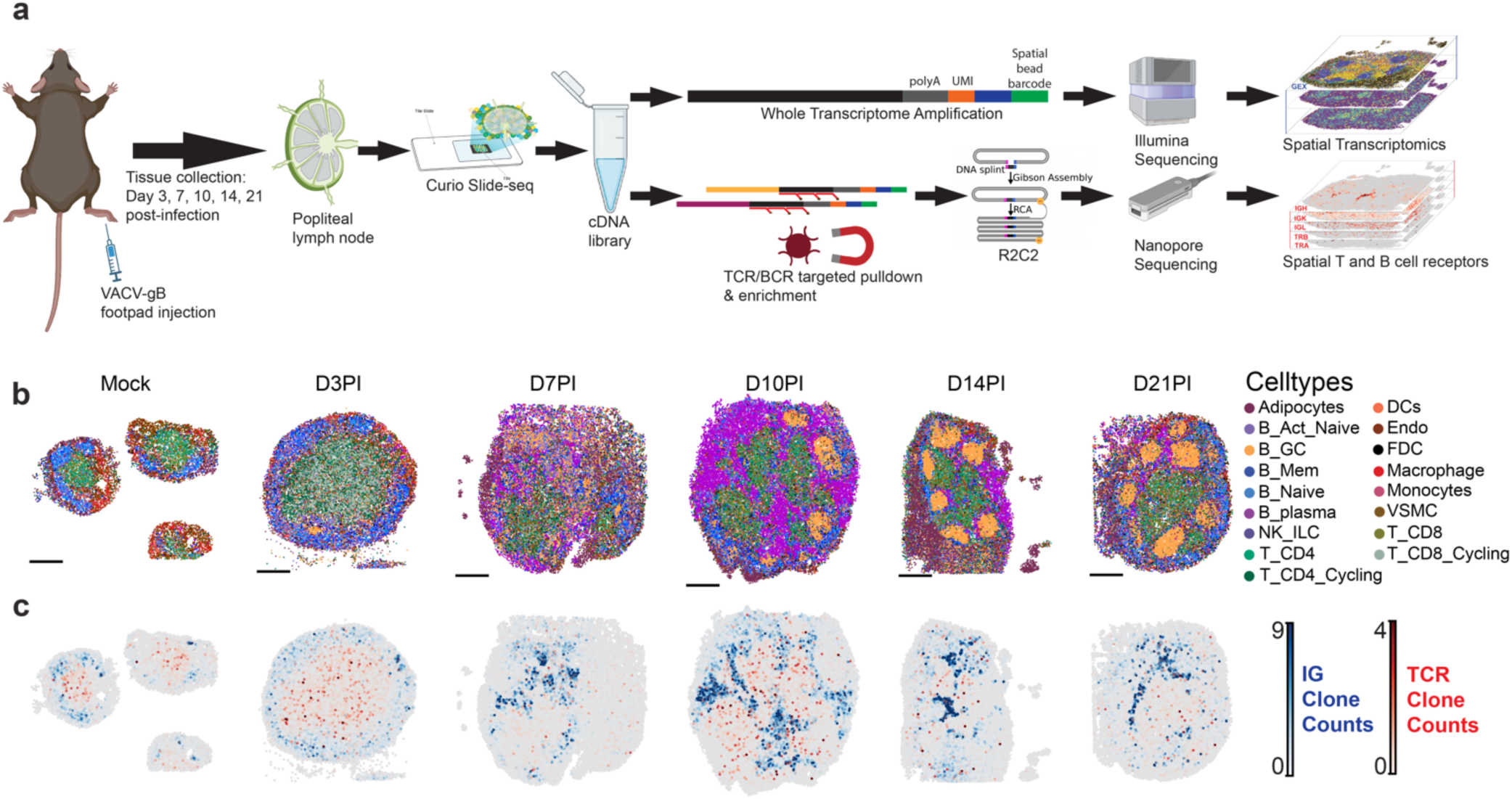
AIR-SPACE enables the mapping of adaptive immune receptor clonotype and transcriptomics *in situ*. **a.** Schematic of the experimental design and methodology, including the generation of long-read (LR) and short-read (SR). **b.** Spatial mapping of cell types across the LN sections at different time points post-infection; scale bars represent 500 μm. **c.** Spatial mapping of adaptive immune receptor (AIR) clonotypes across the LN sections, with immunoglobulin (IG) clones shown in blue and T cell receptor (TCR) clones shown in red.

The AIR-SPACE protocol begins with a 10-µm tissue section mounted onto a spatial transcriptomics array (Curio Seeker, **Methods**). After RNA hybridization, reverse transcription, tissue clearing, second-strand synthesis, and complementary DNA (cDNA) amplification, the resulting cDNA library is split into two portions (**Fig. 1a**). One portion is reserved for short-read sequencing (Illumina), and the other portion is reserved for long-read sequencing of TCR and BCR transcripts (**Fig. 1a**, **Methods**). To specifically enrich BCR and TCR transcripts while retaining spatial barcodes, UMIs, and full-length cDNA, we used hybridization capture with probes tiling the constant region (**Supp. Table 1**). For long-read sequencing, we combined Oxford Nanopore Technology (ONT) with rolling circle to concatemeric consensus amplification (R2C2)^21^ to achieve high-fidelity, full-length sequences, enabling precise characterization of the adaptive immune repertoire.

We employed this assay to study the temporal response of the LN to footpad injection with VACV-gB (**Methods**). We analyzed mouse PLN tissues at 3, 7, 10, 14, and 21 days post-infection (DPI). In addition, we analyzed mock controls collected at day 3 post-injection with PBS (**Fig. 1a**, **Methods**). Spatial transcriptomic sequencing enabled us to resolve the spatial structure of the LN by assigning cell types to beads using deconvolution method with a combined lymphoid organ single-cell RNA sequencing (scRNA-seq) datasets^22–24^ as a reference (**Methods**). Furthermore, by normalizing and aggregating beads into 30 µm bins, followed by integrated clustering, we identified and annotated spatially distinct regions of the LN, including the medulla, outer and inner cortex, germinal centers, and conduit areas (**Methods**).

The spatial arrangement of cell types in our LNs corresponded well with known anatomical organization and reflected the temporal remodeling associated with the infection response. Specifically, T cells were predominantly localized to the paracortex (inner cortex), B cells were concentrated in the outer cortex, and macrophages were primarily distributed within the medulla and capsule regions of the LN. These patterns align well with canonical compartmentalization of LN. Over time post-infection, cycling T cells significantly increased in D3PI compared to Mock, while GC B cells and plasma cells emerged at later timepoints (D10PI, D14PI, D21PI). Additionally, other cell types, including endothelial cells, dendritic cells, and follicular dendritic cells were observed in their expected regions within the LN structure over the infection time course (**Fig. 1b, Supp. Fig. 1e**). Notably, we observed non-random organization of CD4 and CD8 T cells within the paracortex, with CD8 T cells more centrally located and CD4 T cells predominantly at the periphery, which could facilitate effective antigen-specific T cell activation and coordination of adaptive immune responses^8^ (**Supp. Fig. 1a**). We validated cell type assignments, by examining the expression of canonical marker genes (**Supp. Fig. 1b**). To confirm the cell type dynamics captured in the spatial data, we performed fluorescence-activated cell sorting (FACS) analyses on multiple PLNs collected concurrently with those used for spatial transcriptomics. The composition of cell types measured by FACS closely mirrored trends observed in the spatial data. For example, plasma cells expanded rapidly between days 7 and 10 before declining, while T-cells exhibited a sharp increase from day 3 to day 7, followed by a gradual decrease at later time points (**Supp. Fig. 1 c&d**).

### AIR-SPACE reveals the temporal and spatial dynamics of adaptive immune receptor repertoires post-infection

To investigate the adaptive immune response following infection, we acquired an approximate total of 13 million reads by nanopore sequencing, with an average of 2 million reads per sample (**Supp. Table 2**). We obtained high-fidelity consensus reads from the R2C2 reads using C3POa and BC1^25^. The consensus reads were demultiplexed from their putative barcode and UMI sequences, which were extracted from their relative positions anchored by the Curio adapter sequence. This process yielded a total of 990,767 consensus demultiplexed unique reads matching the bead-barcode whitelist, averaging 165,128 reads per sample. We assessed the fidelity of the reads, by evaluating the constant region mapping identity score of immune repertoire sequences using IgBlast^26^. This analysis revealed a median accuracy of 99.7% in the final processed reads (**Fig. 2a**).

**Figure 2.**
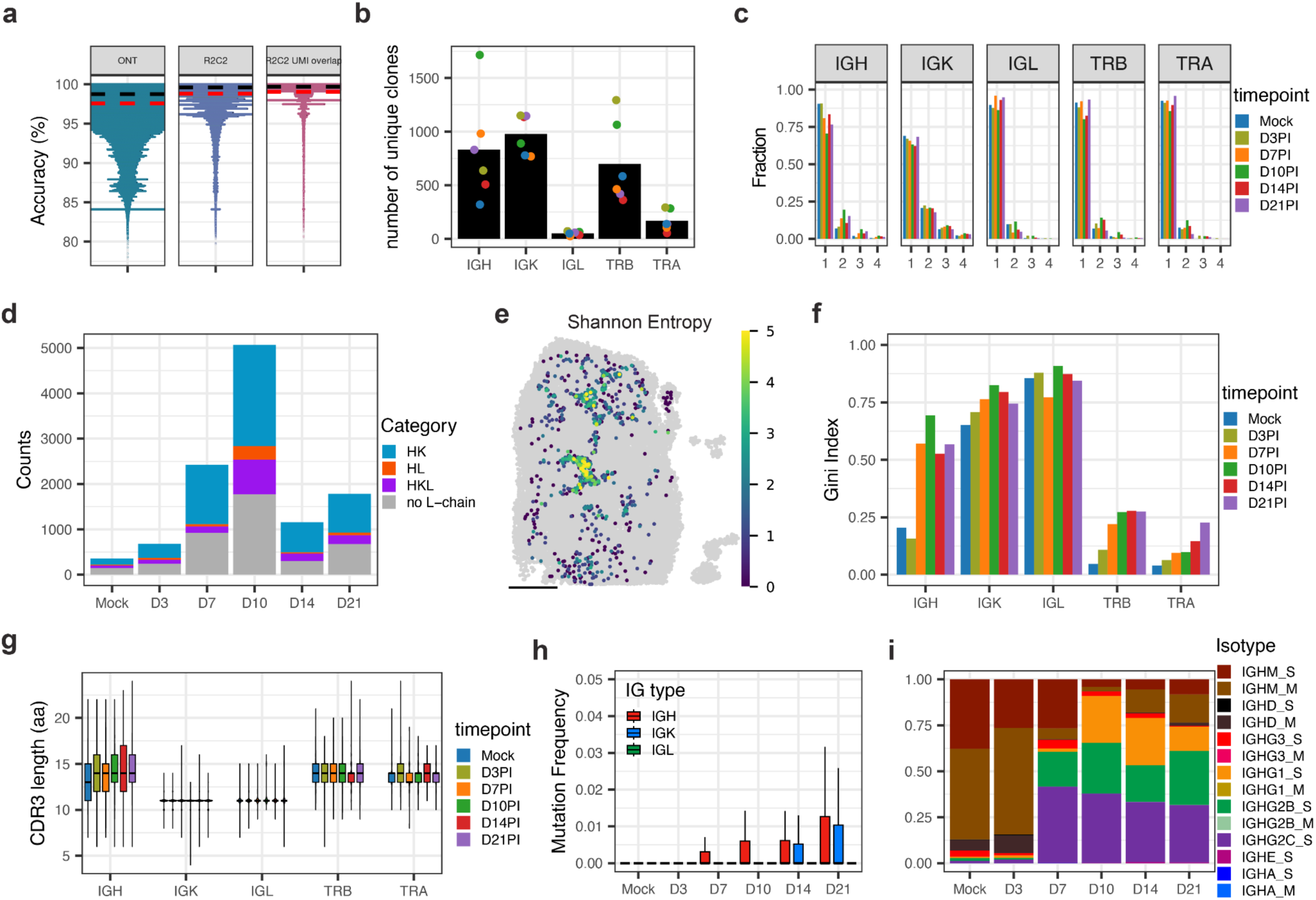
Comprehensive analysis of adaptive immune receptor profiles from long-read sequencing. **a.** Swarm plots comparing the accuracy of ONT reads post-basecalling (ONT), following R2C2 processing (R2C2), and after UMI correction and barcode sequences were mapped to the Curio barcode whitelist (R2C2-UMI overlap). **b.** Barplot displaying the average number of unique clones identified for each receptor type (IGH, IGK, IGL, TRB, TRA) from LRs (R2C2-UMI overlap), with individual sample values represented by dots. **c.** The fraction of beads containing a single or multiple clonotype sequence for each receptor type across samples. **d.** Barplot showing the number of beads with detected heavy chains, categorized by the presence of paired IG receptors (HK: IGH and IGK; HL: IGH and IGL; HKL: IGH and IGK&L; no L-chain: no light chains were detected). **e.** Spatial mapping of Shannon entropy index on LN of D14PI, calculated by potential combinations between heavy and light chains. Scale bars represent 500 μm. **f.** The Gini index represents the diversity of clonotypes for each receptor type across samples. **g.** Boxplot showing the CDR3 length (aa) for each receptor across samples. **h.** Boxplot showing the mutation frequency among IG receptors across samples. **i.** Bar charts show the composition of IGH chain isotypes across samples, with M as the membrane-bound BCR and S as the secreted antibody.

For clonotype annotation of the TCR and BCR transcripts, we used MiXCR^27^ to annotate the V(D)J regions and identify clonotypes within our dataset. The clonotype calling was defined by complementarity determining region 3 (CDR3) sequences sharing the same V and J gene segments while allowing one mismatch or indel within the N-junctional diversity of CDR3 (**Methods**). This analysis identified a total of 6,045 IGH clones, 6,799 IGK clones, 359 IGL clones, 5,287 TRB clones and 1,291 TRA clones (**Fig. 2b**). After demultiplexing and clonotype calling, we mapped the AIR spatially from the long-read data, confirming that BCR-annotated reads predominantly localized to B cells, and TCR-annotated reads were mainly assigned to T cells (**Fig. 1c, Supp. Fig. 2d**). We observed a strong correlation between short-read and long-read data for each clonotype, supporting the robustness of our approach (**Supp. Fig. 2e**).

To assess the spatial resolution achieved by AIR-SPACE for AIR clonotype calling, we examined the clonotype content of individual beads. We found that most beads with annotated clonotype contained a single clonotype sequence across all time points, with slight variations observed between receptor types (**Fig. 2c**). Overall, 83.7% of beads with annotated clonotype contained a single clonotype, including 82.1% of beads with IGH CDR3 sequences and 87.9% of beads with TRB CDR3 sequences, indicating that AIR-SPACE achieves single-cell resolution in AIR annotation for most beads. This aligns well with previous observations from Slide-TCR-seq^19^.

BCRs or antibodies comprise heavy and light chains, whereas TCRs comprise beta and alpha chains. Because paired BCR and TCR sequences determine antigen binding, experimental and computational methods have been developed to identify paired chains within individual B and T cells^28,29^. Using AIR-SPACE, we identified a total of 7,412 beads with paired IG heavy and light chain CDR3 sequences, with an average of 64.5% of beads containing heavy chain CDR3 across samples (**Fig. 2d**). Among these beads, an average of 34% had unique heavy-light chain CDR3 sequence combinations (Shannon entropy value of 0). For those beads without unique combinations, we observed that they are enriched in the conduit region of the LN, likely because of the high density of plasma cells packed in these regions (**Fig. 2e, Supp. Fig. 4b**).

LNs serve as essential secondary lymphoid organs where the initiation and regulation of adaptive immune responses take place^4–6^. We assessed the dynamic temporal changes in the AIR repertoire in the LNs as a function of time post-infection. First, we estimated the diversity of each receptor repertoire using the Gini index, where a value of 1 represents maximal inequality among values. We observed a similar trend of decreasing diversity (increasing Gini index) over time post-infection on all B cell receptors, while T cell receptors remain relatively stable (**Fig. 2f**). Notably, the diversity for the IG light chain repertoires (IGK, IGL) was consistently lower compared to the heavy chain. Additionally, we observed a greater number of shared light chain clonotypes across samples compared to heavy chains (IGH: 2, IGK: 521, IGL: 58). Both of these observations potentially reflect a higher level of coherence among light chains, consistent with previous findings of promiscuous light chains^28^.

For each demultiplexed consensus read that could be clonotype-annotated, we measured CDR3 length and mutation frequency relative to the corresponding germline V(D)J gene (**Fig. 2g**). We also annotated IGH chain isotypes and determined whether they are membrane-bound (BCR) or secreted (antibody). B cells undergo two key processes in LNs: somatic hypermutation, which introduces somatic mutations in the variable regions, and class-switch recombination, which changes the isotype of produced antibodies^30^. Both are mediated by the enzyme activation-induced cytidine deaminase (AID), encoded by the gene *Aicda*. The AIR-SPACE dataset captured expected temporal dynamics within the IG repertoires (IGH, IGK, IGL). IGH CDR3 length progressively increased over time (**Fig. 2g**), and mutation frequencies exhibited a gradual rise over time for both IGH and IGK (**Fig. 2h**). In parallel, IG isotype composition shifted from IgM/D at early timepoints (D3PI, Mock) to predominantly secreted IgG antibodies from D7PI to D21PI. Notably, we observed an increased proportion of IgM/D isotypes and a decreased proportion of IgG3/1 at D14PI and D21PI, suggesting the potential recirculation of naive B cells or the egress of antibody-secreting plasma cells from the lymph node. (**Fig. 2i**).

Next, we examined whether AIR-SPACE could dissect cellular heterogeneity at the individual bead level. We combined the region annotations with IGH clonotype information in a single heatmap (**Fig. 3**). This analysis revealed marked temporal and spatial variations in isotype distribution across the LNs. For example, in the mock and D3PI samples, beads containing IgG isotypes were predominantly located in the medulla, consistent with known LN architecture. As anticipated, the beads in GCs showed higher expression of *Aicda* at later time points (D14PI and D21PI) (**Supp. Fig. 2f**). Interestingly, at D14PI and D21PI, beads with IgM/D isotypes were primarily found in the outer cortex, suggesting the infiltration or recirculation of naïve B cells into these areas. Additionally, we also observed that multiple isotypes within the same bead, especially in the conduit and medulla regions, indicative of dense packing of plasma cells (**Supp. Fig. 2g**). These regions also exhibited a higher rate of somatic hypermutation (**Fig. 3**). Overall, AIR-SPACE resolved fine-scale immunological heterogeneity and dynamic changes in immune cell populations over time.

**Figure 3.**
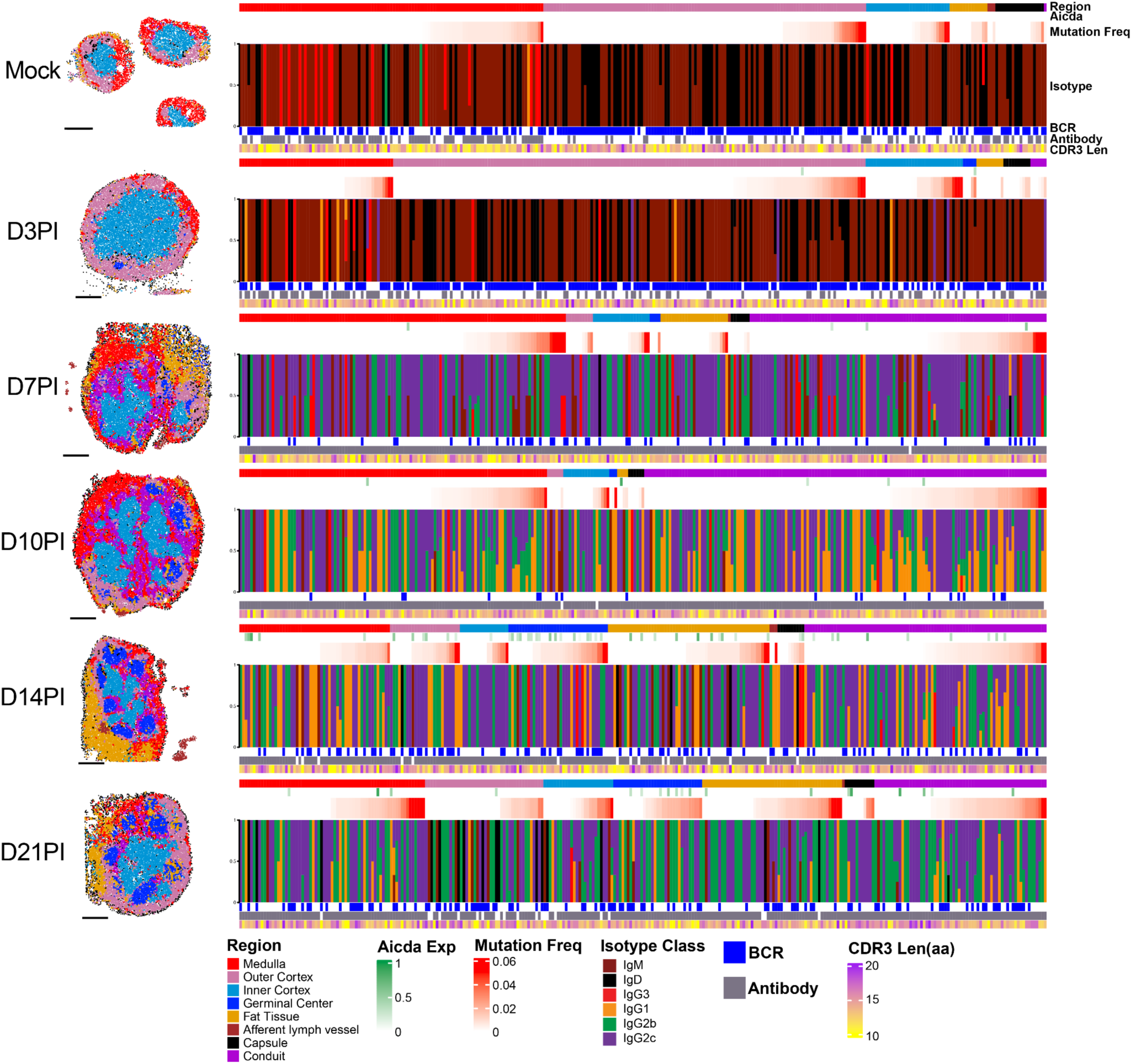
Heatmap of individual beads with IGH clonotype sequences. Left column: Spatial mapping of regions across the LN sections at different time points post-infection; scale bars represent 500 μm; Right column: Heatmap showing individual beads across all samples containing IGH clonotype sequences, mapped to its structural location within the LN (Region), its expression of Aicda, its mutation frequency of IGH, its isotype composition, isoform of BCR or antibody, and its CDR3 length. (To maintain consistent heatmap widths for visualization, the number of beads shown was downsampled to 300, following the same region composition ratios for each sample)

### Spatial niches of interferon-gamma activation in the LN early post-infection

Interferon-gamma (IFN-γ) is a type II interferon that plays an essential role in controlling viral infections by inhibiting viral replication and enhancing both innate and adaptive immune responses^31^. We observed significantly elevated *Ifng* expression at D3PI, followed by a rapid decline at later time points (**Fig. 4a, Supp. Fig 5a**). At D3PI, *Ifng* expression localized to distinct LN regions (**Fig. 4b**). Using K-means clustering, we identified six Ifng-high clusters and categorized beads within a 200 µm radius of their centroids as “Inner niches” or “Outer niches,” corresponding to their location in the inner or outer cortex, respectively. We then quantified *Ifng* expression as a function of distance from these niche centroids. Outer niches showed higher overall *Ifng* expression, and both inner and outer niches showed decreasing *Ifng* levels with increasing distance from their centroids—a gradient absent in a control niche centered away from Ifng-high areas (**Fig. 4c, Methods**).

**Figure 4.**
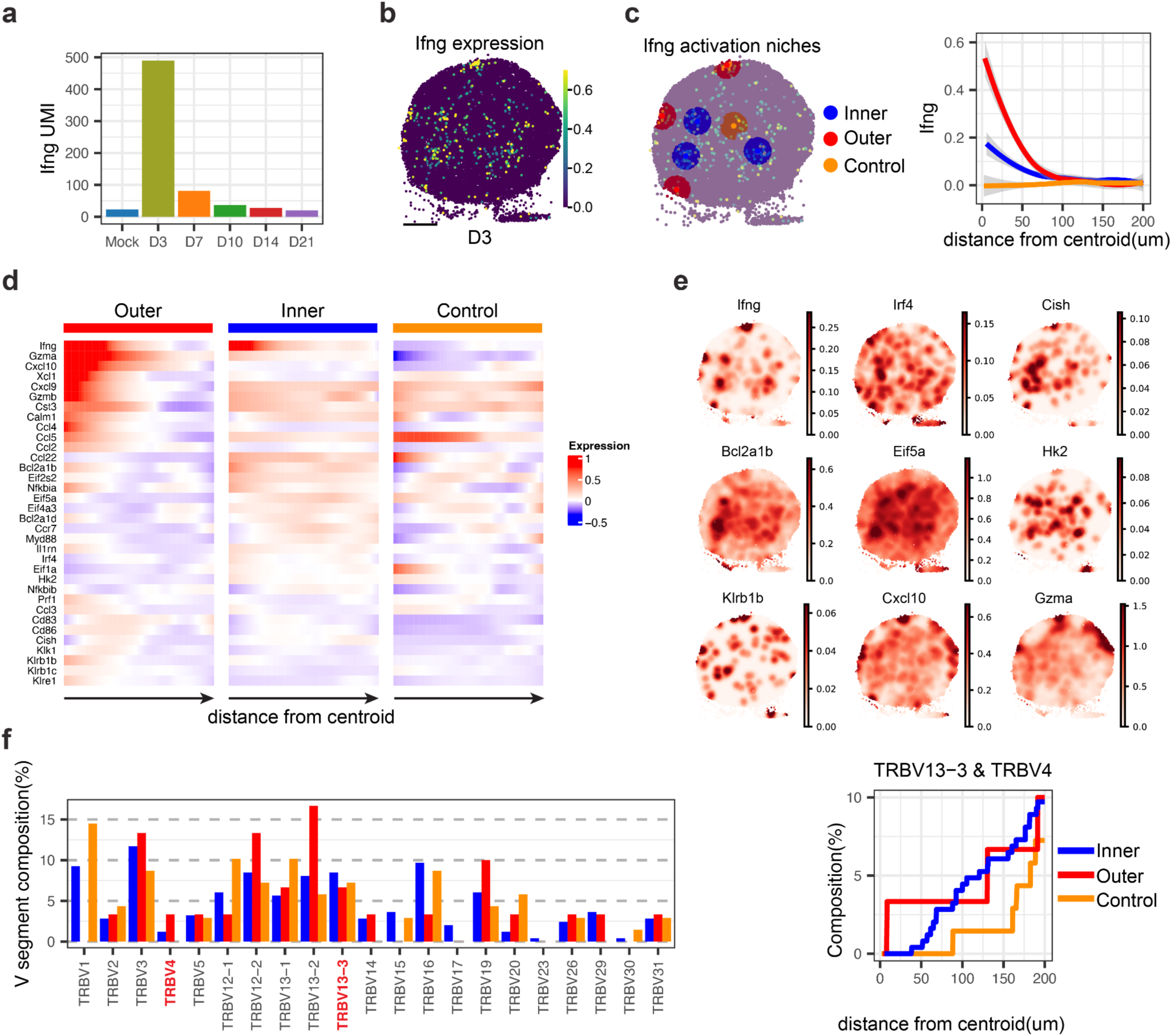
Niches of activation from the expression on Ifng. **a.** Total expression of *Ifng* across all samples at different time points post-infection. **b.** Spatial expression of *Ifng* in D3PI LN. **c.** Left: Identification of Ifng activation niches, categorized as inner (blue), Outer (red), and Control (orange), corresponding to niches in the inner cortex, outer cortex, and central areas with low expression of Ifng in the inner cortex, respectively; Right: *Ifng* expression levels as a function of distance away from the centroid of the three groups of niches, the color was shared with the right panel. **d.** Heatmap showing the expression levels of significantly spatial auto-correlated genes as a function of distance away from the centroid of the niches. **e.** Spatial map on examples of genes positively correlated with *Ifng* in Gaussian smoothing value. **f.** Composition of TRBV gene segments in the three groups of niches, with a detailed focus on TRBV13-3 and TRBV4 composition as a function of distance away from the centroid of niches. The color

Previous studies^32,33^ indicate that NK cells are the primary early producers of IFN-γ prior to the arrival of CD8 T cells, which contribute to later IFN-γ production, providing critical antiviral control. To characterize the transcriptional phenotypes associated with Ifng-rich areas, we performed spatial autocorrelation analysis (**Fig. 4d, Methods**). Gene set enrichment on Ifng-correlated genes (Pearson’s r > 0.4**, Methods, Supp. Table 3**) revealed distinct pathway signatures. Outer niches were enriched in cytokine and chemokine signaling (*Cxcl10*, *Cxcl9*, *Ccl5*) and NK cell-associated genes (*Klrb1b*, *Klrb1c*). We also found high expression of *Prf1*, *Gzmb*, and *Gzma*, suggesting NK cell-mediated cytotoxicity. In contrast, inner niches displayed higher NF-κB signaling (*Nfkb1a*, *Nfkb1b*, *Ikbkb*), Th17 cell differentiation (*Irf4*, *Il1b*), and apoptotic processes (*Bcl2a1d*, *Bcl2a1b*, *Bcl2l1*, **Fig. 4e & Supp. Fig. 5 b&c**). These transcriptional profiles suggest NK cell-driven IFN-γ in the outer cortex and T cell-driven IFN-γ in the inner cortex, in line with previous reports^32,33^. The spatial heterogeneity of *Ifng* expression in the inner cortex suggested antigen-specific T-cell activation. Examining T cell clonality by analyzing TCRβ V segments (TRBV13-3, TRBV4) known to respond to VACV-gB^20^ revealed no significant overall differences among Inner, Outer, and Control niches; however, TRBV13-3 and TRBV4 usage increased linearly with proximity to the centroid in Inner niches—an effect absent in the Control niche (**Fig. 4f**). This finding indicates that antigen-specific T cell responses likely shape localized IFN-γ expression patterns within the LN inner cortex.

### AIR-SPACE characterizes germinal center compartments and B cell spatial dynamics

GCs are specialized microanatomical sites in secondary lymphoid organs where B cells undergo clonal expansion and antibody affinity maturation^30^. To investigate the spatial organization and dynamic evolution of B cells within GCs, we focused on samples from the later time points (D10PI, D14PI, D21PI) when GCs were mature and well-developed (**Supp. Fig. 6a**). Among them, two GCs stood out as they exhibited clearer microanatomical structures (**Fig. 5 a&b**). In the D14PI GC1, unsupervised clustering with GraphST^34^ identified Light and Dark Zones (LZ and DZ, **Supp. Fig. 6b**). LZ markers (*Cxcl13* and *Cxcr5*) were more highly expressed in the LZ, with expression levels increasing progressively as beads were positioned farther from the LZ-DZ boundary. Conversely, DZ markers (*Cxcl12*, *Cxcr4*) formed corresponding spatial gradients (**Fig. 5c**, **Supp. Fig. 6d**). These findings align with the well-established DZ/LZ molecular architecture, driven by chemokines (*Cxcl12*, *Cxcl13*) and their receptors^30^ (*Cxcr4*, *Cxcr5*). Differential expression analysis further highlighted functional differences between LZ and DZ (**Supp. Fig. 6c**). *Aicda* (p = 3.3e-07) and G2M scores (p < 2.22e-16) were elevated in DZ, consistent with active clonal expansion and somatic hypermutation. *Bcl2a1b* (p = 0.0039) and the FDC marker *Mfge8* (p = 3.8e-12; PMID: 18490487) were higher in the LZ, suggesting antigen-driven selection. We observed very similar patterns in another GC in D21PI LN, indicating polarized LZ and DZ subregions (**Fig. 5d**).

**Figure 5.**
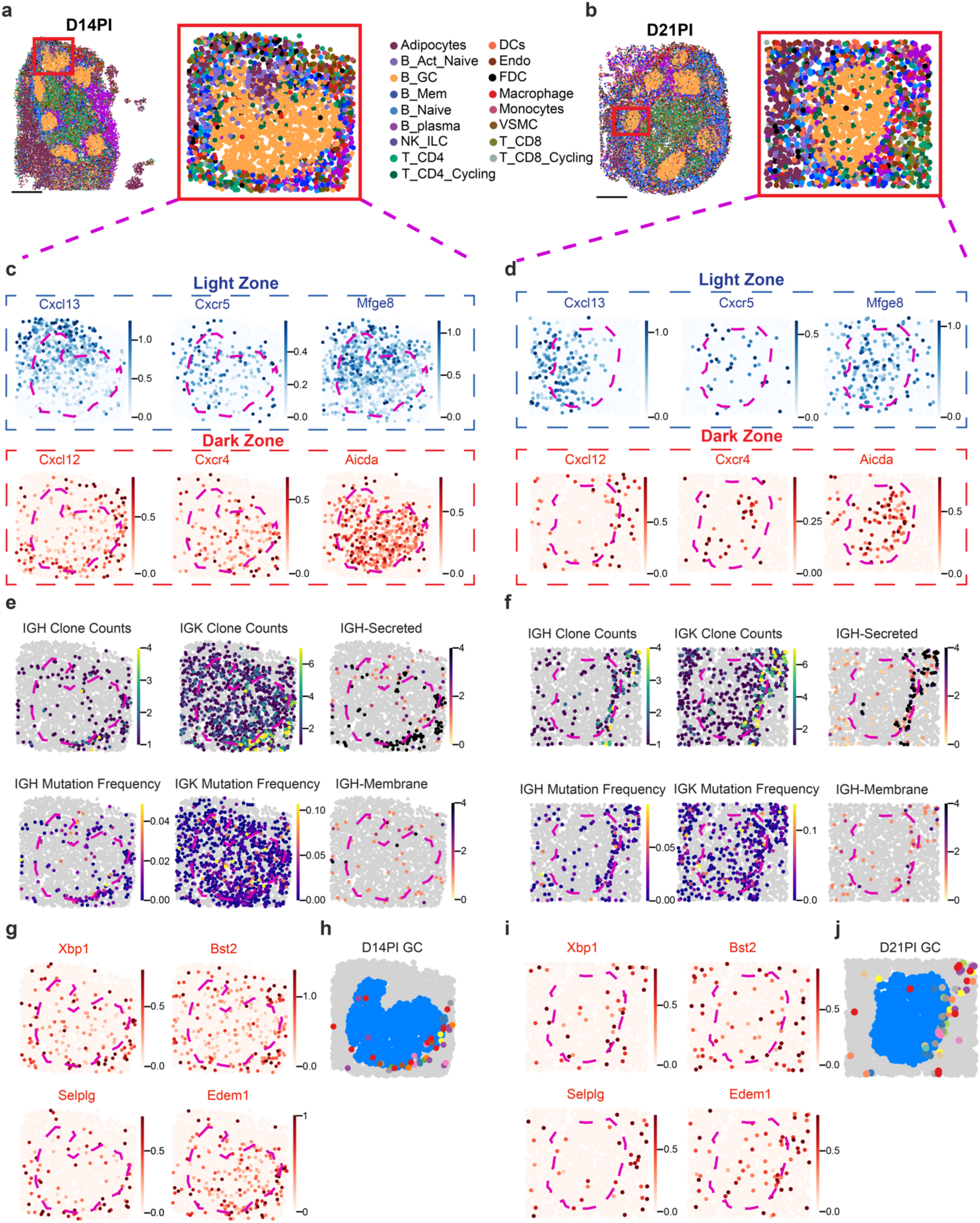
AIR-SPACE uncovers dynamic changes in germinal centers. a&b. Zoomed-in view of one GC from both the D14PI and D21PI samples, colored by different cell types; Spatial mapping of cell types across the LN sections at different time points post-infection; scale bars represent 500 μm. **c&d.** Spatial expression of LZ (*Cxcl13, Cxcr5, Mfge8*) and DZ (*Cxcl12, Cxcr4, Aicda*) marker genes. **e&f.** Spatial mapping of adaptive immune repertoire information from LR. **g&i.** Spatial expression of pre-plasma cell marker genes (*Xbp1, Bst2, Selplg, Edem1*). **h&j.** Spatial maps of the spatial location of different IGH clones on the GCs from D14PI and D21PI, color-coded by different IGH clones.

Strikingly, we identified numerous beads at the DZ–medulla interface displaying plasma cell–like features: high *Xbp1* expression, low *Aicda* expression, abundant IGH secreted isoforms, and elevated IGH mutation frequency (**Fig. 5e&f, Supp. Fig. 6c**). By applying spatial autocorrelation analysis on these GCs and investigating gene expression highly correlated with IGH secretion processes (**Supp. Table 4**), we identified several putative pre-plasma cell markers^10^, including *Bst2* and *Selplg* (**Fig. 5g&i**). We also found genes associated with “Protein processing in the endoplasmic reticulum” (*Mzb1*, *Edem1*), “N-Glycan biosynthesis” (*Man1b1*, *Ddost*), and “Antigen processing and presentation” (*H13*, *H2-Q7*) (**Supp. Fig. 6i&j**). These pathways are characterized as features of plasma cell differentiation due to their roles in supporting the production of high quantities of antibodies^35^. Importantly, by spatial tracking identical IGH clones within the GC, we observed that these cells appear to migrate out of the GC via the DZ interface (**Fig. 5h&j**). These findings support the theoretical model^9^ that these cells are pre-plasma cells exiting the GC through the DZ after selection and differentiation. Collectively, these findings demonstrate the utility of AIR-SPACE to capture both the transcriptional differences and the clonal dynamics of B cells, including the emergence and egress of pre-plasma cells, thereby providing new insights into the spatial and temporal complexity of B cell-plasma cell differentiation within GCs.

Recent studies in mouse and human LNs demonstrate that individual B cell clones can expand and undergo selection in multiple GCs, indicating B cell recirculation^14,36^. To investigate this phenomenon through the lens of AIR-SPACE, we performed the spatial lineage analysis and examined the spatial distribution of IGH clones across GCs. We first assigned each IGH clone (with at least five beads in the spatial data) to a GC by measuring the Euclidean distance between their bead coordinates and the identified GCs (**Supp. Fig. 6a**). Beads within 100 µm were assigned to that GC. If multiple beads from the same clone were associated with different GCs, the clone was categorized as “Multiple GCs.” Clones with all beads assigned to the same GC or with no GC assignment were categorized as “Single or non-GC” clones. Applying this classification to LNs with developed GCs (D10PI, D14PI, D21PI) showed that although most IGH clonal families were confined to a single GC, a notable fraction spanned multiple GCs (**Fig. 6a**). Interestingly, the proportion of “Multiple GCs” clones increased over time: 6.9% at D10PI, 17.5% at D14PI, and 22.9% at D21PI (**Fig. 6a**). The number of clones assigned to each GC range from 35 to 261 with median 86 clones per GC. In agreement of previous study^36^, the distribution of the number of shared IGH clones by the number of GCs fits well with a Poisson distribution, indicating that the recirculation of a B cell to a different GC is a stochastic mechanism (**Fig. 6b**). The spatial distribution of the inter-GC clones show they located across multiple GCs (**Fig. 6c**). Taken together, these findings support the idea of recirculation of clones and inter-GC exchange as has been suggested previously by others^36^.

**Figure 6.**
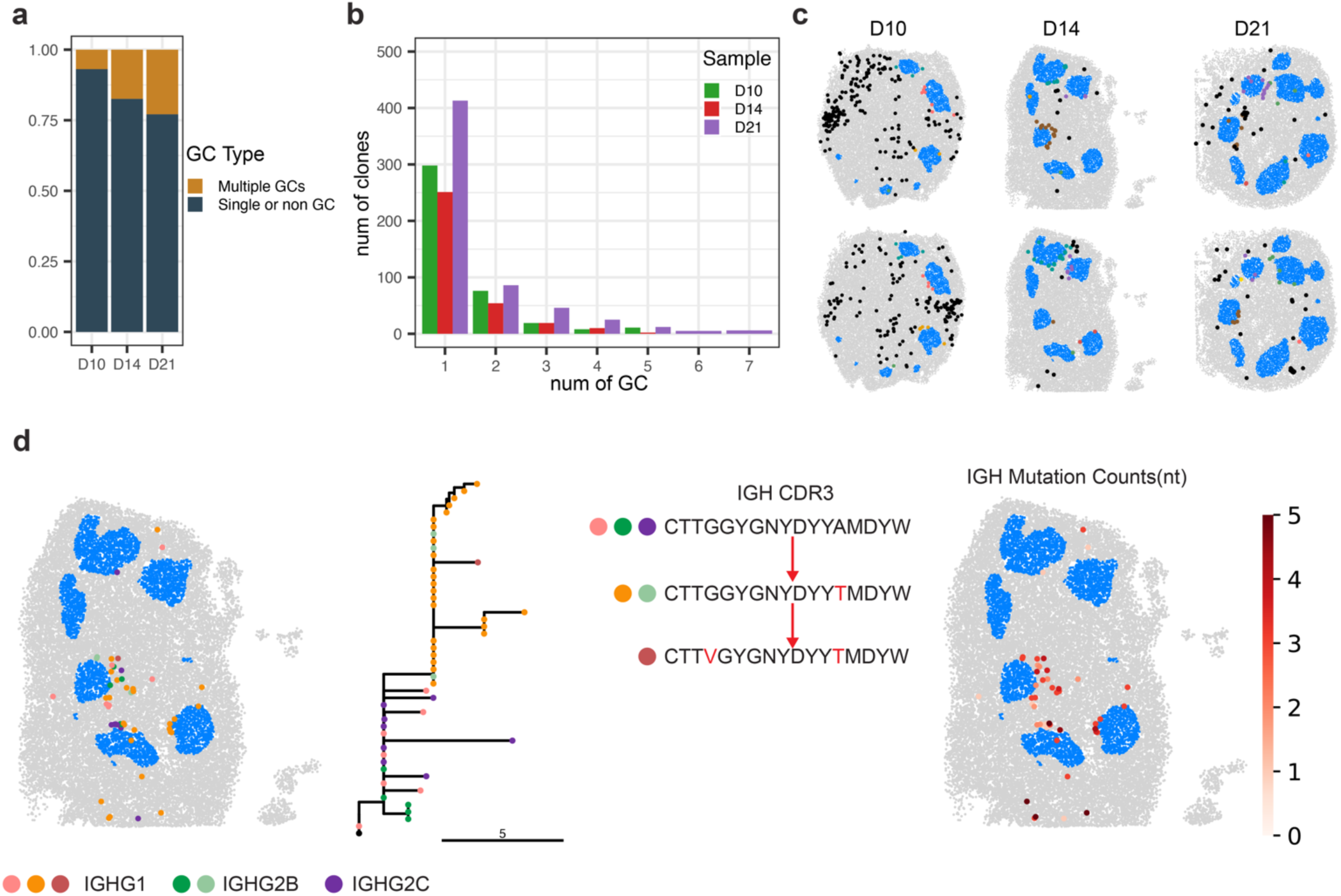
AIR-SPACE uncovers dynamic changes in germinal centers. **a.** Percentage of IGH clones found in multiple GCs (brown) or single GC (black) at different time points (D10PI, D14PI, and D21PI). **b.** The distribution of the number of IGH clones by the number of GCs for each sample. **c.** Spatial map showing examples of IGH clones shared across multiple GCs for different time points (D10PI, D14PI, and D21PI). Blue dots represent individual GCs, black dots indicate beads not assigned to any GC, while colored dots correspond to beads assigned to adjacent GCs. **d.** Clonal evolution of a single IGH clone family from the D14PI sample. Spatial mapping and lineage tree represent each individual bead within the family, colored by different isoforms and CDR3 sequences. Branch lengths reflect the number of mutations (nt) accumulated from the germline (black node: root), and its spatial map.

Finally, to explore sequence-level analysis of immune cell clonal evolution, we constructed phylogenetic trees for a subset of prominent IGH clonal families and investigated their lineage diversification and mutation accumulation as a function of location within the LN (**Methods**). One D14PI IGH clone stood out: The lineage tree of this clonal family revealed a branching structure indicative of ongoing somatic hypermutation and class switching recombination (**Fig. 6d**). Its variants were spatially distributed across 3 neighboring GCs, as well as regions not assigned to any GC in the outer cortex of the LN. These spatial patterns imply that B cells shuttle among multiple GCs as they undergo affinity maturation. Together, these data indicate that GCs are not isolated microenvironments. Instead, they appear to function as interconnected nodes in a dynamic network, allowing B cells to circulate among multiple GCs, potentially enhancing the efficiency and diversity of the affinity maturation process.

## DISCUSSION

In this study, we introduce AIR-SPACE, a method that integrates high-fidelity long-read sequencing of target-enriched adaptive immune receptor transcripts with unbiased spatial RNA sequencing. We used this method to construct an atlas of the temporal and spatial dynamics of immune repertoire changes in mouse popliteal lymph node (LN) tissues over the course of VACV-gB infection. The immune cell clonotypes measured in this way localized to their expected LN structures, and their temporal changes aligned with the established trajectory of the adaptive immune response. The data further provide new insights into the spatiotemporal dynamics of the LN microenvironments following viral infection.

First, we examined mechanisms of T cell priming in the LN during the early response to infection. Previous work^32,33^ has shown that IFN-γ is essential for antiviral immunity, with NK cells providing early production to enhance innate responses, while CD8 T cells sustain IFN-γ levels during the adaptive phase to suppress viral replication and prevent tissue damage. However, the reprogramming pathways of the IFN-γ activation and the spatial components within the microenvironment remain poorly defined. By mapping IFN-γ expression, we identified discrete activation niches in the early post-infection LN that exhibited heterogeneous gene expression profiles depending on LN location. Niches in the outer cortex were marked by strong innate immune signatures, driven by NK cells and macrophages, whereas niches in the inner cortex displayed gene expression patterns indicative of T cell activation, likely involving antigen-specific T cells as corroborated by *in situ* TCR clonotype data. It is known that virions can reach the draining LN within hours of infection, triggering rapid innate IFN-γ production and CD8 T cells priming by dendritic cells^5,7,37^. We anticipate that utilizing AIR-SPACE to study these very early dynamics will yield novel perspectives into these mechanisms.

AIR-SPACE also enabled in-depth characterization of the spatial organization of GC microenvironments and B cell spatial dynamics within the LNs. Notably, we gained deeper understandings into the differentiation of GC B cells to plasma cells. Meyer-Hermann et al.^9^ previously used computational modeling to study these differentiation dynamics and concluded that B cells likely leave the GC as plasma cells via the DZ rather than the LZ. While prior classical models^38,39^ assumed that GC B cells leave the GC directly via the LZ after selection. In this model^9^, after positive selection in the LZ, GC B cells migrate back to the DZ, where they undergo division and differentiate into plasma cells if their antigen was retained, and ultimately exit the GC via the DZ. This model is supported by imaging and mRNA microarray analysis^11,12^ which furthermore highlighted the role of T follicular helper cells in driving plasma cell egress from the DZ. Consistent with the model and these studies^9–12^, we identified a clear population of plasma cells around the DZ–medulla interface with high expression of *Bst2*, *Selplg*, and *Xbp1*, low expression of DZ B cell markers (*Aicda* and G2M proliferation score), and upregulation of endoplasmic reticulum and protein export–related genes. Furthermore, by analyzing the positioning of B cell clonotypes across the DZ–medulla interface, we observed a gradient increase at this interface in antibody expression and IGH mutation frequency—clear plasma cell features. Although our data offer only snapshots of this dynamic process, the distribution of identical IGH clones around the DZ–medulla interface provides valuable clues regarding the GC B-to-plasma cell egress pathway. Collectively, our observations provide molecular and genetic evidence that antibody-secreting plasma cells exit the GC via the DZ–medulla boundary.

A deeper understanding of the mechanisms governing the development and dynamics of B cell clones within and between GCs is crucial for vaccine design^14^. Tas et al.^14^ used multiphoton imaging and a multicolor confetti mouse system to reveal sharing of B cell clones between GCs within the same LN^14^. Later, Pelissier et al.^36^ used a combination of laser capture microdissection of individual GCs and repertoire sequencing to substantiate the evidence for B cell recirculation among GCs in LNs^36^. Here, by mapping B cell clonotypes spatially using AIR-SPACE, we likewise observed shared B cell clones across multiple GCs. Moreover, our data offer a significant advantage by enabling spatial lineage tracing of a B cell clonal family within a single LN section via phylogenetic tree analysis, allowing us to better understand the heterogeneity of these dynamic processes not only among different GCs but also across various B cell clones. Applying AIR-SPACE to study antigen boosting may illuminate secondary responses that involve the formation and reactivation of memory B cell clones^40^.

AIR-SPACE offers several advantages over previous spatial antigen receptor methods. First, a key advantage of is its ability to characterize both T and B cell clonotypes *in situ* at high 10-μm spatial resolution, allowing to resolve paired TCR/BCR sequences within the same spot. Second, AIR-SPACE is compatible with multiple commercially available platforms for spatial transcriptomics and does not require specialized equipment, making it easily adoptable by others. Looking ahead, AIR-SPACE opens new avenues for exploring adaptive immune responses in a variety of contexts, for example, investigation into immune responses to various pathogens^41,42^, autoimmune conditions^43^, gut microbiome-immune relationships^44,45^, or tertiary lymphoid structures within cancers^46^. Moreover, integrating AIR-SPACE with emerging spatial total transcriptome methods^45,47^ or spatial proteomics technology^48^ could broaden the scope for studying systems immunology. As spatial sequencing technologies continue to evolve, enhancements in resolution, sensitivity, and scalability will likely expand the utility of AIR-SPACE, enabling more comprehensive insights into the cellular and molecular landscapes of immunity. Ultimately, the knowledge gained from such advancements have the potential to guide novel antibody design strategies and deepen our understanding of immune-mediated diseases.

## DATA AVAILABILITY

Data will be made available upon publication under GEO accession number: GSE286452.

## CODE AVAILABILITY

Code associated with this work can be found at https://github.com/ShaowenJCornell/AIR-SPACE

## ACKNOWLEDGMENTS

We thank the Cornell Biotechnology Resource Center for their help with sequencing the libraries. We thank the Cornell Center for Animal Resources and Education for animal housing and care. We thank Lena Takayasu, Rohit Agarwal, and other members of the De Vlaminck lab for helpful discussions and feedback. This work was supported by NIH R01AI176681, NIH R01AI176681, and CZI 2023-323354 to I.D.V. and R01AI189855 to I.D.V. and B.R.

## AUTHOR CONTRIBUTIONS

SJ, MM, BDR, and IDV conceived of the study. SJ, MM, and PS performed the spatial transcriptomics and downstream experiments. VM and SB performed the animal experiments. SAL performed the FACS experiments. SJ performed the data analysis. SJ and IDV wrote the manuscript. All authors provided feedback and comments.

## METHODS

### Sample information and processing

Mature adult male mice (3-month-old; C57BL/6J; male) were injected with 1e5 PFU Vaccinia virus (VACV-gB) in the left hind footpad via 30G needles. Equal volume of PBS was injected to the mice of the same age as controls. These virus-infected mice and controls across five different time points, namely D3 (infected and Mock), D7, D10, D14, D21, were used for spatial transcriptomics experiments. After isoflurane anesthesia, the left PLNs of these mice were extracted using an aseptic technique and were immediately embedded in cryomolds with O. C. T Compound (Tissue-Tek) media and fresh frozen directly on dry ice and stored at -80 °C.

### Flow cytometric processing and analyses

Single cell suspensions were prepared from lymph nodes and were washed with 1X PBS. Cells were then stained with eBioscience Fixable Viability Dye eFluor 780 (ThermoFisher Scientific) or Ghost Dye Violet 510 (Cytek) for 30 min at 4 °C. Next, cells were washed in Fluorescence-Activated Cell Sorting (FACS) buffer (1X PBS with 2% normal calf serum) and incubated for 10 min in anti-CD16/CD32 (clone 2.4G2, Bio X Cell) diluted in FACS buffer. Mixtures of fluorochrome-conjugated antibodies to stain surface markers were diluted in FACS buffer and then used to stain cells for 1 h at 4 °C. Antibodies used to stain surface markers included: anti-mouse CD4 (BUV805, clone GK1.5, BD Biosciences), anti-mouse CD4 (BV650, clone RM4-5, BD Biosciences), anti-mouse CD4 (BUV395, clone GK1.5, BD Biosciences), anti-mouse CD19 (R718, clone 1D3, BD Biosciences), anti-mouse CD8a (BUV395, clone 53-6.7, BD Biosciences), anti-mouse Sca-1 (BV605, clone D7, BD Biosciences), anti-mouse TACI (Alexa Fluor 647, clone 8F10, BD Biosciences), anti-mouse CD138 (PE-Cy7, clone 281-2, Biolegend), anti-mouse IgD (APC-H7, clone 11-26c.2a, BD Biosciences), anti-mouse GL7 (PE, clone GL7, BD Biosciences), and anti-mouse FAS (PE-CF594, clone Jo2, BD Biosciences). Cells were then washed with FACS buffer. Cell suspensions were then fixed and permeabilized with the BD Pharmingen Transcription Factor Buffer Set (BD Biosciences) according to manufacturer’s instructions. When BCL6 staining was required, permeabilized cells were stained for 45 min at 4°C with anti-mouse BCL6 (BV711, clone K112-91, BD Biosciences) diluted in 1X Perm/Wash solution (BD Biosciences). Cells were washed with FACS buffer and resuspend in FACS buffer for flow cytometry acquisition. Flow cytometry samples were acquired on a BD FACSymphony A3 Cell Analyzer (BD Biosciences) and analyzed with BD FACSDiva V9.0 and FlowJo V10.10 software (BD Biosciences).

### Slide-seq spatial transcriptomics library preparation and sequencing

Slide-seq spatial transcriptomics experiment was performed using the Curio Seeker Kit (Curio Bioscience) according to manufacturer instructions. In brief, a 10-µm thickness tissue section from each collected fresh-frozen PLN was mounted on a 3-mm x 3-mm spatially indexed bead surface Curio Seeker tile. After RNA hybridization and reverse transcription, the tissue section was digested, and the beads were removed from the glass tile and resuspended. Second-strand synthesis was then performed by semi-random priming followed by cDNA amplification. A sequencing library was then prepared using the Nextera XT DNA sample preparation kit (Illumina). The library was sequenced on an Illumina NextSeq 2K (Illumina) platform using a P3 100bp kit, with reads allocated as follows: 50 bp for read 1, 8 bp for index 1, 8 bp for index 2, and 72 bp for read 2.

### Histological processing

For hematoxylin & eosin (H&E) staining, we used the serial 10-µm thickness sister sections from the section used from the spatial transcriptomics. These sections are fixed in pre-chilled methanol for 30 min and then processed following the 10x Visium H&E staining protocol. H&E stained PLN tissue sections were imaged using a Zeiss Axio Observer Z1 microscope equipped with a Zeiss Axiocam 305 color camera. The resulting H&E images were corrected for shading, stitched, rotated, thresholded, and exported using Zen 3.1 software (Blue edition).

### Hybridization-based target enrichment of TCR and BCR transcripts

Hybridization-based target enrichment of TCR and BCR transcripts was performed using the IDT xGen NGS target enrichment kit. BCR and TCR custom-designed 5’-biotinylated oligonucleotide panels were used separately for capture and pulldown of target molecules of interest. The BCR panel includes 151 probes consisting of *Igh, Igk, and Igl* gene loci, and the TCR panel includes 81 probes consisting of *Trb, Tra, Trg, and Trd* gene loci (**Supp. Table 1**). In order to have sufficient material, the input cDNA libraries were amplified prior to the hybridization reaction. We used 5-10 ng of Slide-seq cDNA per library. PCR reactions were performed using KAPA HiFi HotStart ReadyMix (2x) (Roche) with cDNA primers (10x Genomics). The total volume and number of PCR cycles of reaction varied depending on the original cDNA amount in each library. After PCR amplification, each library was performed bead wash with 0.6X SPRIselect and eluted in 40 μl of water. Next, we followed the protocol “xGen hybridization capture of DNA libraries” (version 7, IDT) with “Tube protocol” with slight modifications. Specifically, we separated the hybridization reaction for each sample into TCR and BCR individually, using 200 ng of PCR-amplified DNA as the input library for each. Post-capture PCR was performed for each pull-down library using KAPA HiFi HotStart PCR with 14 cycles, followed by two rounds of 0.6X SPRIselect as post-capture PCR clean-up and eluted in 22 μl H_2_O.

### Rolling circle amplification to concatemeric consensus (R2C2)

We first generated the splint by using 23 μl of H_2_O, 25 μl of KAPA HiFi HotStart ReadyMix (2x), 1 μl of UMI_Splint_Forward (100 μM) and 1 μl of UMI_Splint_Reverse (100 μM). The mix was incubated for 3 min at 95 °C, for 1 min at 98 °C, for 1 min at 62 °C and for 6 min at 72 °C. The DNA splint was then purified with the Select-a-size DNA Clean and Concentrator kit (Zymo Research) with 85 μl of 100% EtOH in 500 μl of DNA binding buffer. Next, 200 ng of target-enriched DNA was mixed with 200 ng of DNA splint and 10 μl NEBuilder HiFi DNA Assembly Master Mix (2x) (New England Biolabs, NEB) was added. The mix was incubated for 1 hour at 50 °C. After incubation, each reaction was added 5 μl of NEBuffer 2, 3 μl of Exonuclease I, and 3 μl of Exonuclease III, and 3 μl of Lambda Exonuclease (all NEB), and adjusted the volume to 50 μl with H_2_O. The mixture was then incubated for 6 h at 37 °C followed by a heat inactivation step for 20 min at 80 °C. The circulated DNA was then extracted using 0.8X SPRIselect and eluted in 40 μl H_2_O. After purification, each circularized DNA was split into 4 aliquots of 10 μl. Each aliquot was amplified in its own 50 μl Rolling circle amplification reaction (formula as in R2C2 paper^21^) and incubated at 30 °C overnight. After incubation, T7 Endonuclease was added to each reaction and then incubated for 2 h at 37 °C with occasional agitation. Cleanups with SPRI beads in 0.5x ratio of H_2_O were performed to extract the debranched DNA and eluted in 40 μl H_2_O.

### Oxford nanopore long-read library preparation and sequencing

Each resulting DNA from R2C2 was sequenced using one separate ONT MinION (R10.4.1) flow cell. For each run, 1-2 μg of DNA was prepared using Ligation Sequencing Kit V14 following the manufacturer’s protocol as in “DNA repair and end-prep” and “Adapter ligation and clean-up”. Each library was then loaded into the flow cell according to the protocol instructions. Each run lasted at least 48 h, and the sequencing results were stored in the state-of-art format.

### Long-read data preprocessing

After ONT long-read sequencing, each resulting data was base called using the super accuracy model (SUP) of the GPU accelerated Guppy algorithm (v6.5.7, config file: dna_r10.4.1_e8.2_400bps_sup.cfg). Basecalled reads were then processed and demultiplexed into R2C2 consensus reads and their subreads using C3POa (v3) with corresponding splint sequences (**Supp. Table 5**). After that, BC1 commands were applied in the above reads to generate R2C2+UMI consensus reads^25^. Next, R2C2+UMI reads were demultiplexed in a similar manner as BLAZE pipeline with modification. Specifically, we changed the following parameters in the config.py file from BLAZE^34^ so that the pipeline can work for Slide-seq library structure (ADPT_SEQ=’TCTTCAGCGTTCCCGAGA’; PLY_T_LEN=8; PLY_T_NT_AFT_ADPT=(13,50); DEFAULT_UMI_SIZE= 7; DEFAULT_BC2_SIZE= 6; DEFAULT_BC1_SIZE= 8). After anchoring the adaptor (linker) sequence with an allowance of one Levenshtein edit distance and identifying its position in each read, we extracted the putative barcode and UMI sequences based on their relative position from the adaptor. Next, the resulting reads that can be anchored to the Slide-seq adaptor were further demultiplexed by overlapping the putative Barcode to the corresponding bead barcode location file (whitelist) with an allowance of one Levenshtein edit distance. After demultiplexing, we trimmed the adaptor, bead barcode and UMI sequences for each read and wrote the reads to a new fastq file.

### Immune clonotype data downstream analysis

#### Adaptive immune receptor clonotype calling and analysis

To call the TCR/BCR clonotype, we applied MiXCR (v4.6.0)^27^ with a modified “generic-ont” preset that can assemble clonotypes within one mismatch or indel nucleotide in CDR3 region. The reads that did not map to any clone (cloneId = -1) or could not be aligned to the whitelist were filtered out. We then used the MiXCR output tabular file to assign reads to clonotypes and spatial bead coordination with the matched barcode. Finally, two tabular files were generated from this process: one is a metadata table for each bead barcode to display its clonotype annotation and abundance level reflected by the number of long-read UMI, which can be concatenated to the Anndata object; the other is a table containing each read along with its clonotype information and spatial barcode coordinates.

#### Adaptive immune receptor isotype and mutation exporting

To extract the mutations relative to germline sequence or isotype usage from the clonotype calling results in a tabular form, we applied MiXCR “exportAlignments” function as follows: “mixcr exportAlignments -f -descresR1 -cloneId - allNMutations -allNMutationsCount -nMutationsRate VRegion -chains -isotype subclass -target sequences -isProductive VRegion –impute-germline-on-export -allNLength”.

To resolve whether each IGH transcript encodes a membrane-bound or secreted isoform, we employed a custom R script leveraging local pairwise alignments against a reference of known mouse IGH constant loci. This reference was assembled from IMGT-derived *Mus musculus* IGH constant-region nucleotide sequences, encompassing both membrane-bound (M) and secreted (S) variants for each locus. After the local pairwise alignments, each demultiplexed long-read was assigned to the most closely matching M or S IGH isoforms or “not assigned” based on its highest alignment score and whether the aligned region differed by no more than 3 nucleotides from the reference.

#### Spatial clonal evolution analysis

This analysis is mostly done using the Immcantation framework. To begin this analysis, we first assign VDJ genes using IgBLAST^26^ from Immcantation Lab Docker image. Then we focused on specific IGH clones that are abundant existing in the spatial dataset by extracting relevant long reads with custom settings. We filter out the unproductive sequences, prioritize full VDJ reads, and select the highest-quality alignment if multiple reads share the same CDR3 and barcode. The resulting data frames were merged with the metadata so that each barcode would have only one representative IGH long-read. To facilitate lineage tracing, we used “createGermlines()” function from Dowser^49^ package with IMGT mouse IGH VDJ segments as reference. Observed mutation frequencies across the variable region were computed with “observedMutations()” from shazam^50^ package. The lineage tree construction was done with igphyml^51^, scaling branch lengths by mutation counts. The resulting trees were visualized with plotTrees(), tips were labeled and color-coded by CDR3 sequences and isotype.

### Short-read sequencing analysis

#### Slide-seq data preprocessing for quality control and smear removal

After Illumina sequencing, the raw sequencing reads were aligned to the mouse genome (assembly: GRCm38) using the STARsolo (v2.7.10a)^52^ pipeline to generate a gene x bead barcode expression count matrix. By integrating the corresponding bead barcode location files downloaded from Curio Bioscience website, slide-seq count matrix with spatial position information for each sample was generated and loaded into an AnnData object using Scanpy (v1.9.8)^53^. Beads with less than 100 transcripts captured were filtered. To remove the smear effect, we also calculated the spatial distances between all pairs of beads within each sample and beads with less than 15 beads within 100 um distance were removed.

#### Multilevel cell-type label assignment for spatial transcriptomics datasets

All samples of spatial transcriptomics post-filtered datasets were concatenated to a big Anndata object for performing cell-type label assignment together. First, cell2location^22^ (v0.1.3) was used to deconvolve our spatial transcriptomics datasets with the combined human lymphoid organs scRNA-seq datasets as a reference same as the cell2location paper. To match our mouse data, the single-cell reference^22^ was first converted to mouse gene symbols by using mousipy (v0.1.6). Next, genes in the single-cell reference were filtered with the following parameter (cell_count_cutoff=5, cell_percentage_cutoff2=0.03, nonz_mean_cutoff=1.12) to select the highly-variable-genes and cell type signatures were estimated using a Negative binomial regression model with “Sample” as batch_key and “Method” as categorical_covariate_keys. Then, spatial mapping was performed on our concatenated Slide-seq datasets with this reference model under the following hyperparameters (N_cells_per_location=1 and detection_alpha=20). A further round of clustering was performed on the cell abundance matrix estimated by cell2location to assign celltype with the most abundant label. Furthermore, another round of deconvolution to refine the T cell and B cell subtypes using two different scRNA-seq references (T cell: LN dataset from DestVI paper^23^; B cell: Human Tonsil Cell Atlas^24^) respectively, with the same parameters as above. Additionally, the concatenated object was applied with Harmony across all samples with further Leiden clustering with resolution=1.0. Adipocytes were annotated on above clusters that show Adipocyte-related markers (*Fabp4*, *Cfd*). All cell-type labels were integrated together for visualization and downstream analysis on the concatenated Anndata object via Scanpy package.

#### Bin normalization and region label assignment

First, we applied bin normalization to all sample Slide-seq post-filtered Anndata objects by transforming unit feature size from 10 µm to 30 µm squares, which would help identify the structural anatomical region of the LNs. After binning, we concatenated all samples’ binned Anndata objects to a combined Anndata. Next, log-normalization, highly variable gene identification (min-disp = 0.20, max_mean = 5), scaling and regressing on “total_counts” were performed on this combined Anndata. Dimension reduction was conducted using Principal Component Analysis (PCA), followed by Harmony integration (n_pcs=20) and Leiden clustering (resolution=0.7). Each cluster was annotated to each region based on their marker genes and spatial location together. The region labels were subsequently transferred to unbinned Anndata objects according to the corresponding units within each binned square.

#### Identification of *Ifng* Activation Niches in the D3PI sample Spatial Map

To establish the Ifng activation niches, we first selected bins (30 µm) with high Ifng expression levels. The binned Anndata object from D3PI sample was performed log-normalization, regression, and scaling. Bins with Ifng expression levels greater than or equal to a predefined cutoff (≥ 4) were classified as “Ifng-high”. The spatial coordinates of these Ifng-high bins were used for K-means clustering with the number of clusters set to 6. Based on their spatial location in either the Inner Cortex or Outer Cortex, they were labeled as “Inner” or “Outer”, respectively. The centroids of these clusters were calculated to represent each niche. To establish a “Control” for comparison, we calculated the centroid of the areas that were at least 200 µm away from the “Ifng-high” areas. Beads within a 200 µm radius of the calculated centroids were identified and used to categorize the Ifng activation niches into three categories: “Inner”, “Outer”, and “Control”.

#### Spatial autocorrelation Analysis

To find the top spatially correlated genes with Ifng, we used the computational method Smoothie^54^. The method uses Gaussian smoothing to address the noise and sparsity in the spatial gene expression data and then performs efficient pairwise Pearson R correlation between genes to rank gene pairs from having correlated to anti-correlated spatial patterns. We used a Gaussian standard deviation of 100 µm in this analysis. To identify pathways and gene sets associated with genes positively correlated with *Ifng*, we performed gene set enrichment analysis using the GSEApy (v1.1.2)^55^ package with Enrichr API. Genes with a Pearson correlation coefficient r greater than or equal to 0.4 were selected, and Ifng itself was excluded for enrichment analysis. The resulting list of genes was analyzed using two specific gene sets: “KEGG_2019_Mouse” and “GO_Biological_Process_2023”. The organism parameter was set to “Mouse” to ensure compatibility with the selected gene sets.

