## Supplemental Figures for "A Temporal and Spatial Atlas of Adaptive Immune Responses in the Lymph Node Following Viral Infection"

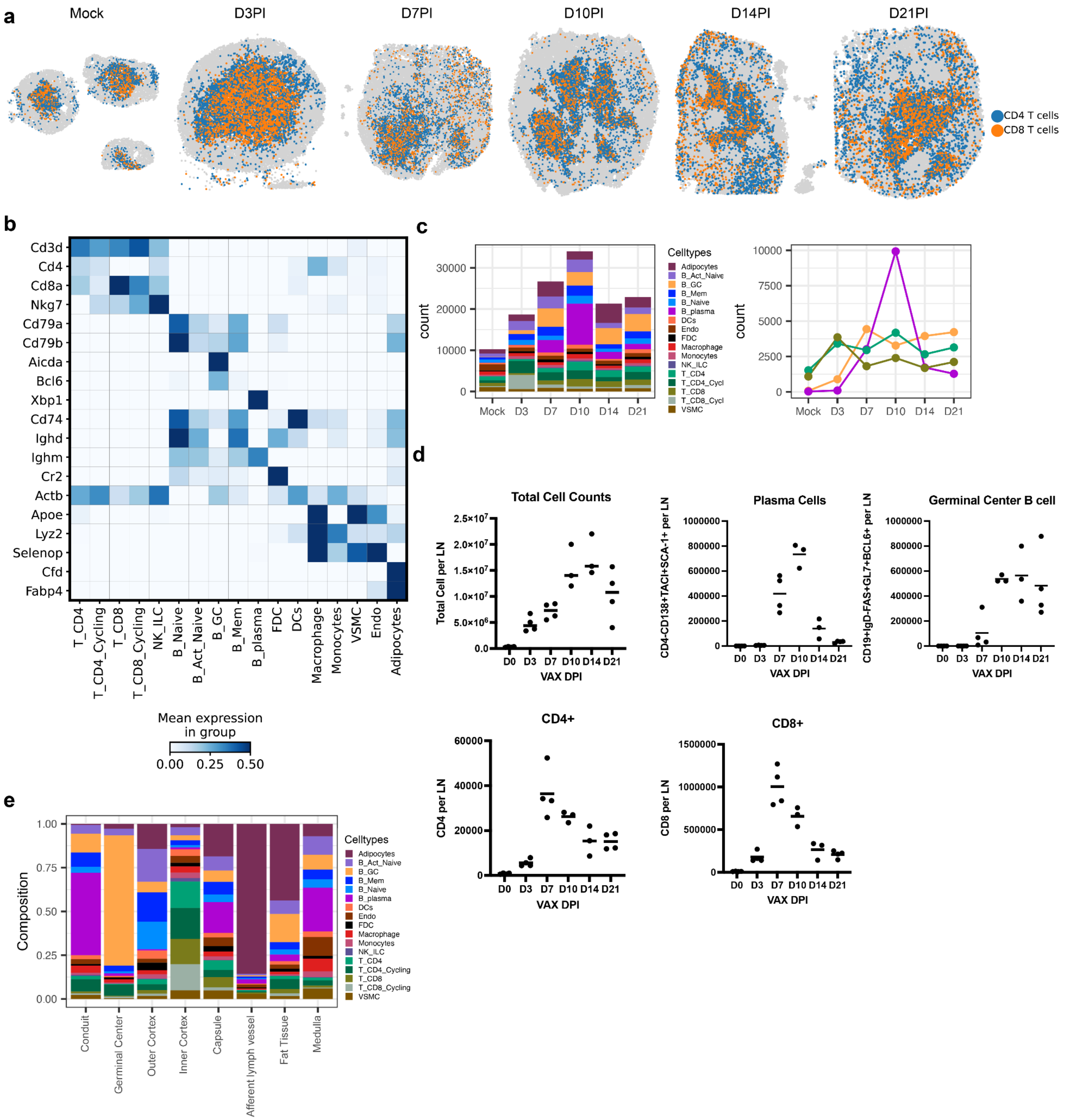


**Supplementary Figure 1. Cell type classification and abundance. a.** Spatial maps show the spatial distribution of annotated CD4/CD8 T cells across all samples. **b.** Matrix plot shows mean expression of cell type marker genes combined with all samples. **c.** Left: Stacked bar charts show total and individual cell type abundance across all samples. Right: Line plots highlighting the selected major cell type abundance (plasma cells, germinal center B cells, CD4 T cells, and CD8 T cells) over the course of infection. **d.** FACS results show total cell counts for LNs at each time point, including plasma cells, germinal center B cells, CD4+ and CD8+ T cells. **e.** Barplot shows the composition of cell types that contributed to each regions within LN, combined all samples.

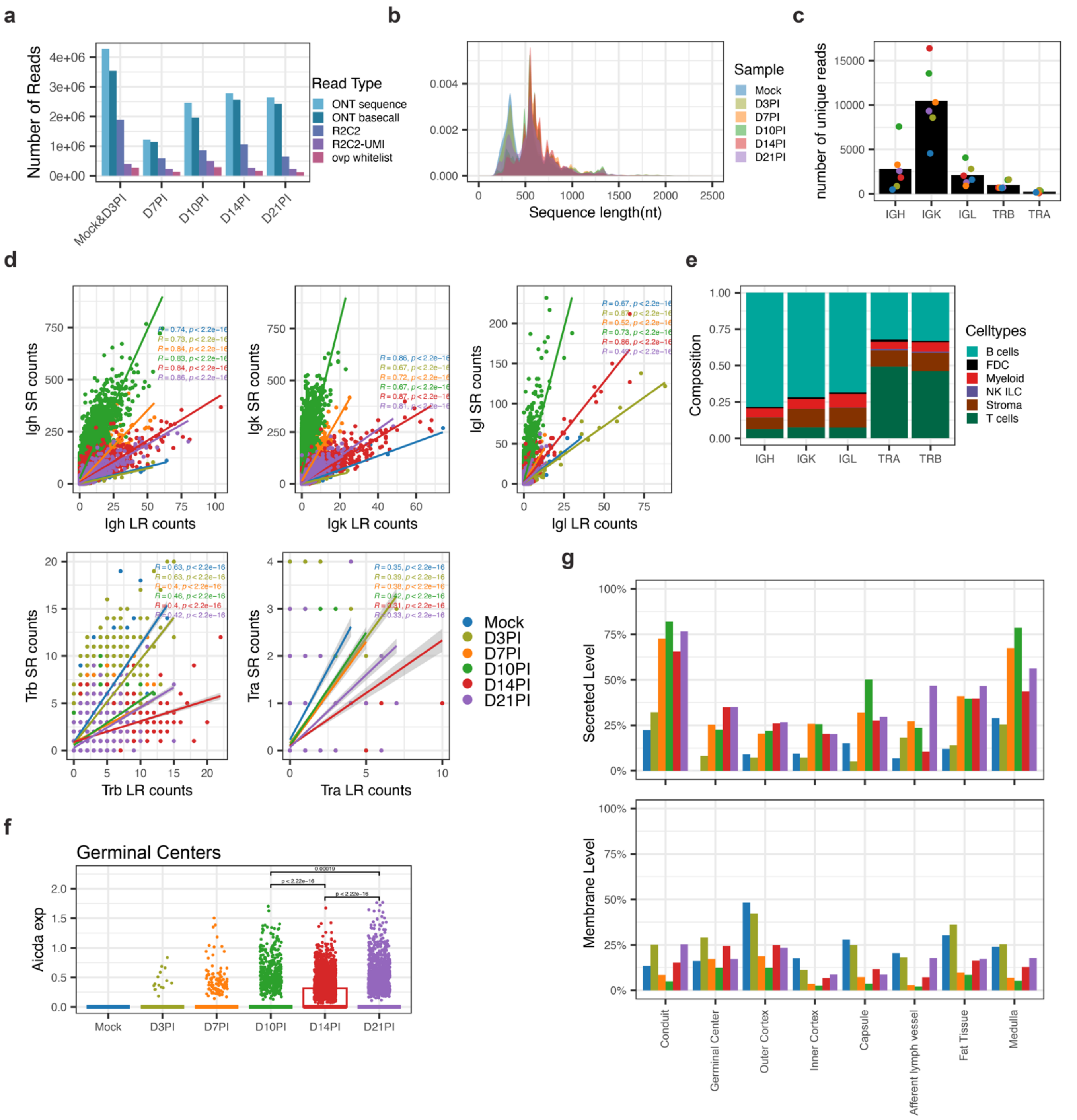


**Supplementary Figure 2. a.** Barplot shows the number of reads obtained at each step of the bioinformatics processing pipeline, originating from ONT LRs to the final consensus reads mapped to the spatial map across all samples. **b.** Distribution of the sequence lengths from the final mapped consensus reads, measured in nucleotide (nt) across all samples. **c.** Barplot shows the average number of unique reads annotated for each receptor type from final mapped consensus reads, with individual sample values represented by dots. **d.** Correlation plots between LR and SR counts for each receptor type across all samples. **e.** Barplot shows the composition of cell types that contributed to the annotated LRs for each receptor type, combined all samples. **f.** Boxplot shows the normalized expression of *Aicda* in all cells within germinal centers across all LNs g. Barplot shows the fraction of beads expressed BCR (Membrane) or antibody (Secreted) in different regions within different samples.

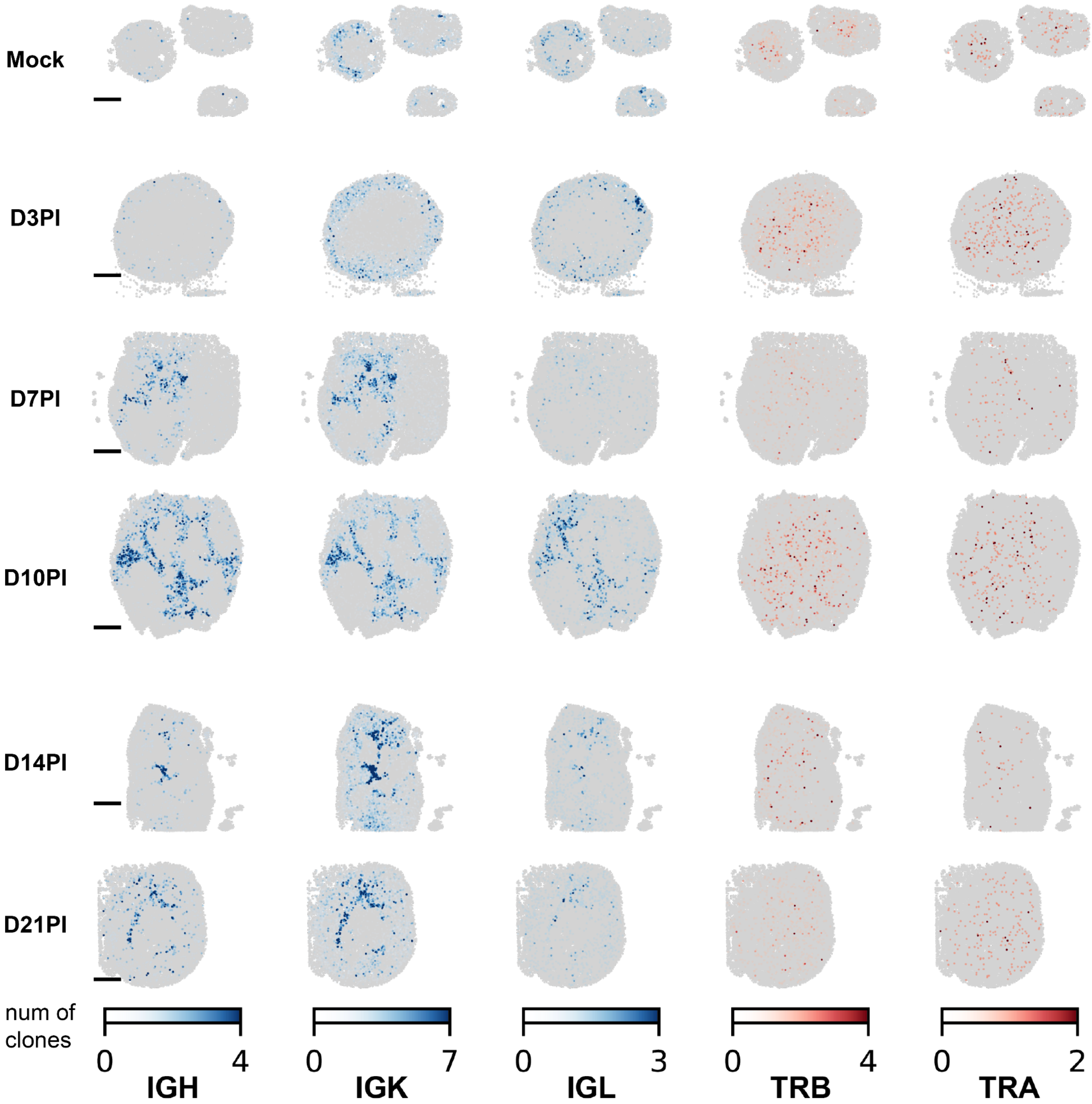


**Supplementary Figure 3.** Spatial mapping of each adaptive immune receptor clonotype across all LN sections, as IGH, IGK, and IGL as shown in blue, TRB and TRA as shown in red.

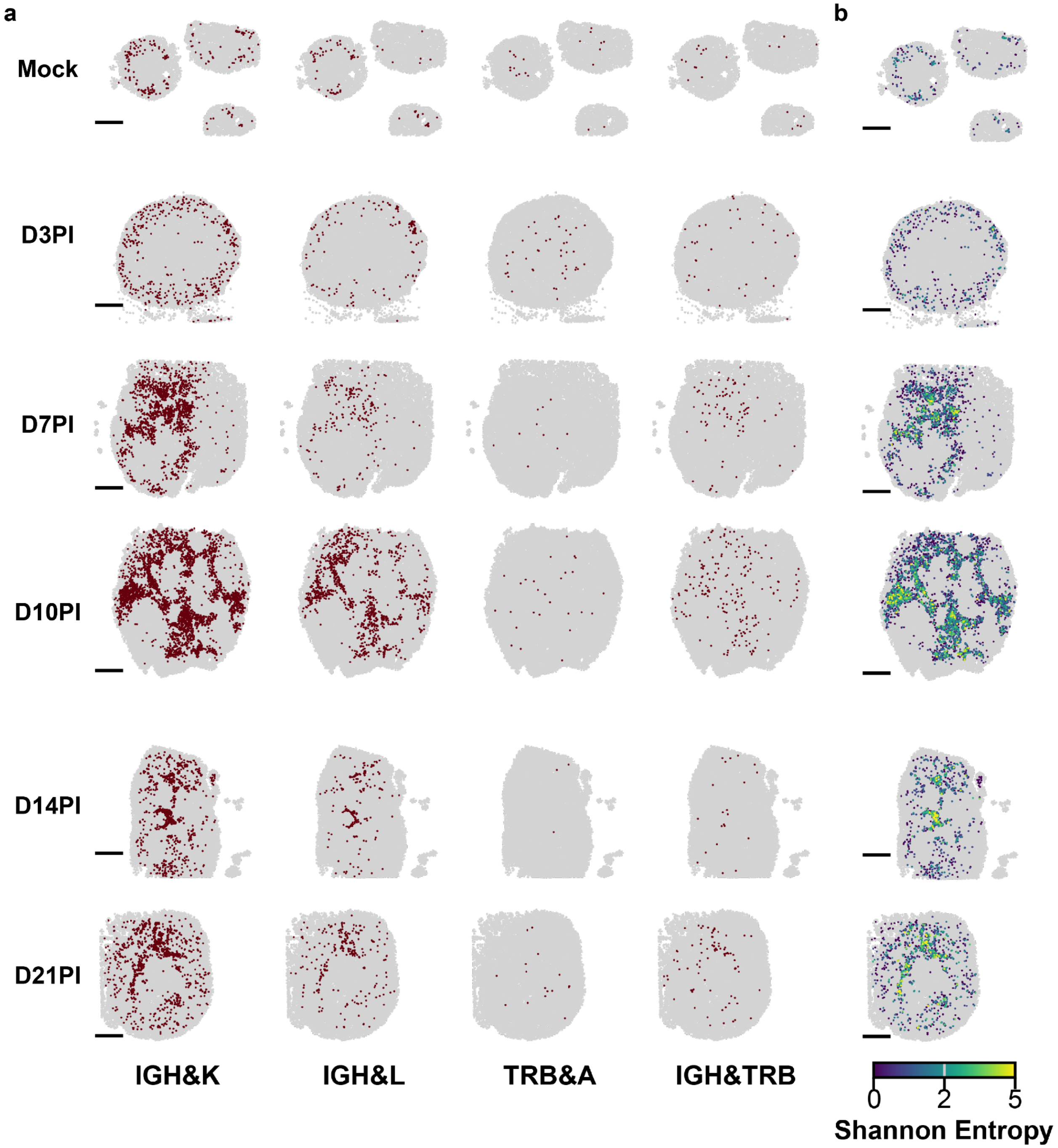


**Supplementary Figure 4. a.** Spatial maps show beads with paired adaptive immune receptors across all LN sections, as paired IGH&IGK, IGH&IGL, TRB&TRA, and IGH&TRB from left to right. **b.** Spatial maps show Shannon entropy value across all LN sections, calculated by potential combinations between IG heavy and light chains.

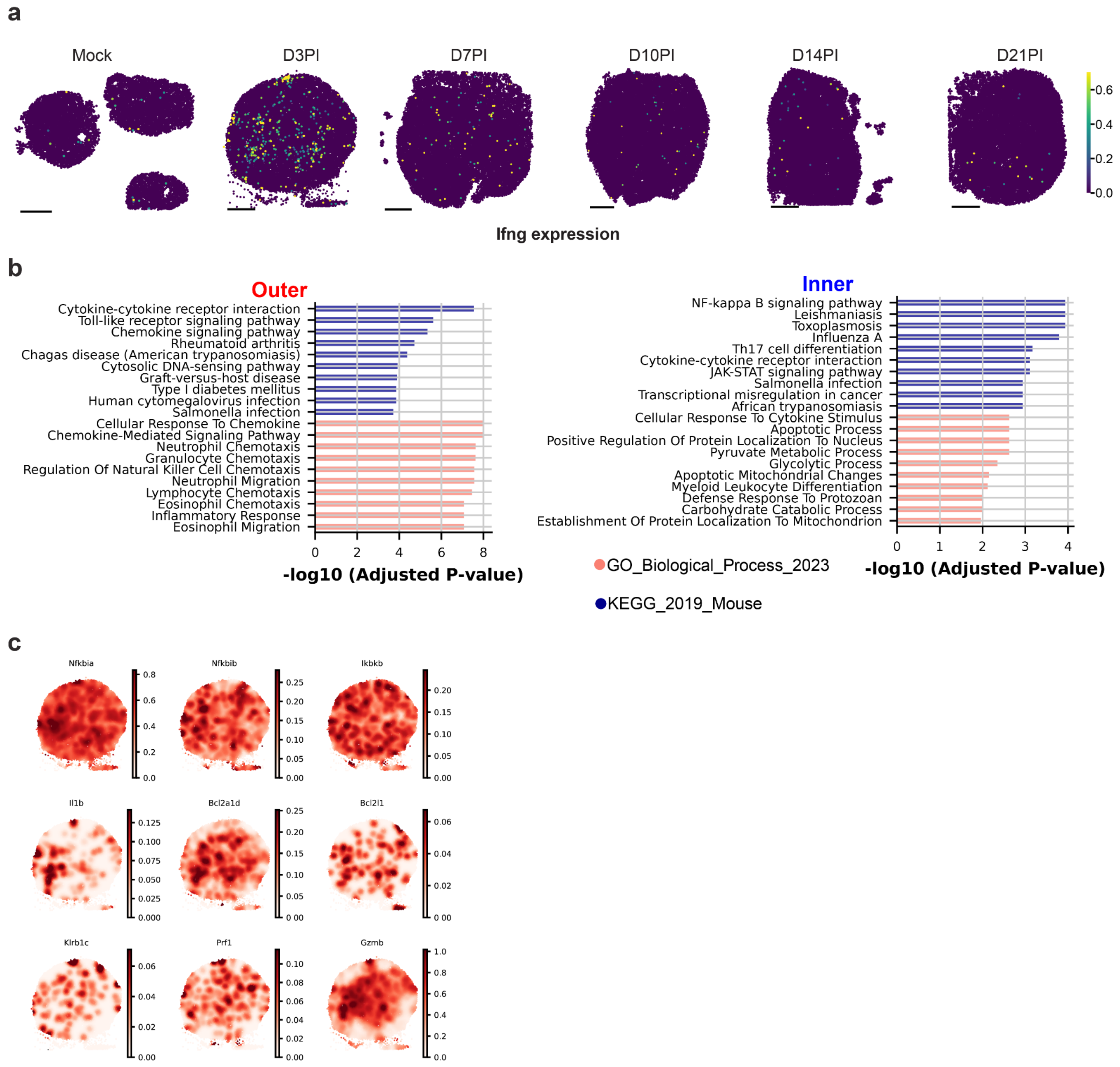


**Supplementary Figure 5. a.** Spatial maps show expression of *Ifng* across all LN sections **b.** GSEA results for genes positively correlated with *Ifng* (Pearson's r > 0.4), Outer and Inner niches respectively. **c.** Spatial map on examples of genes positively correlated with *Ifng* (*Nfkbia, Nfkbib, Ikbkb, Il1b, Bcl2a1d, Bcl2l1, Klrb1c, Prf1, Gzmb*) in Gaussian smoothing value.

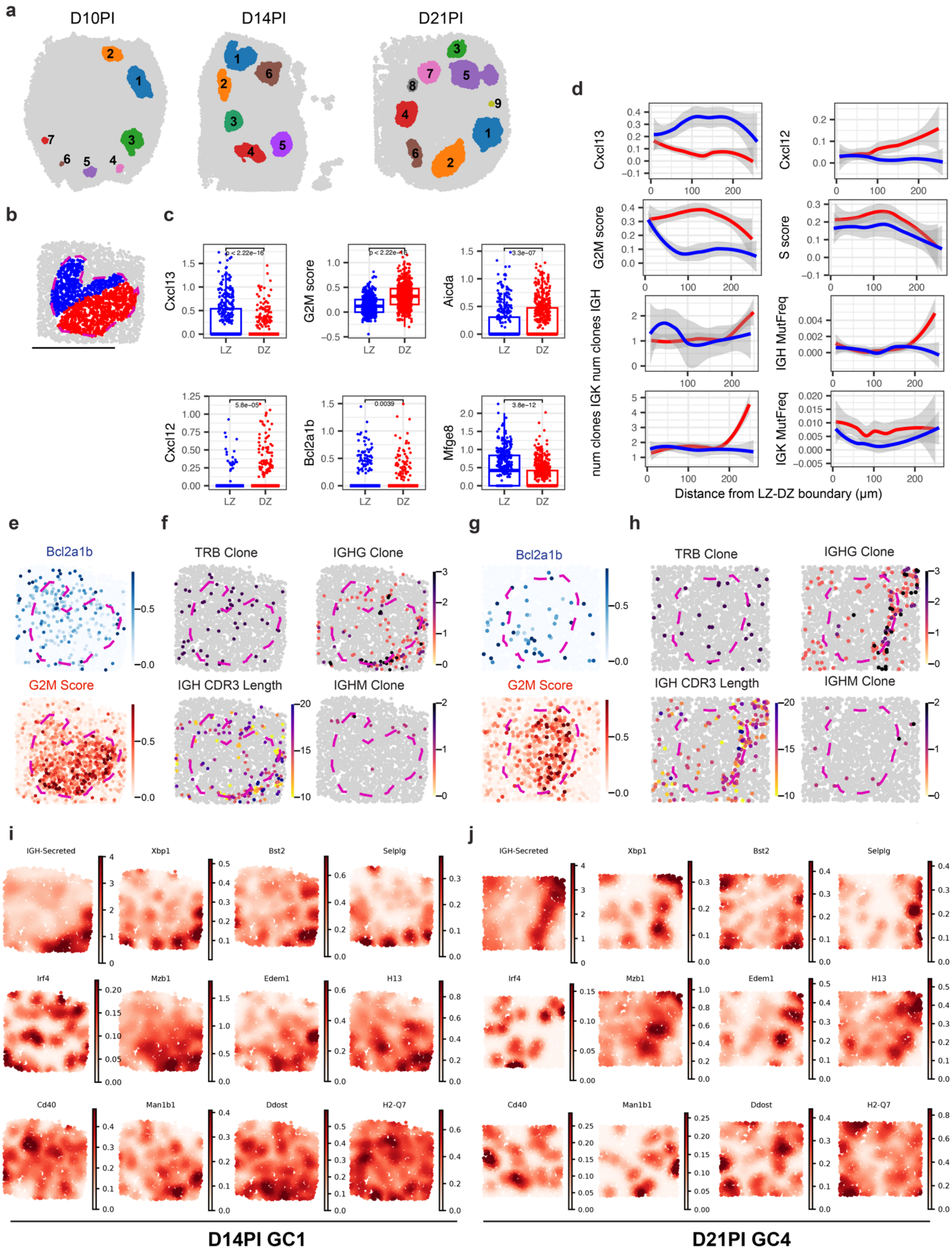
a
D10PI
D14PI
D21PI
b
c
Cxcl13
G2M score
Aicda
Cxcl12
Bcl2a1b
Mfge8
LZ
DZ
d
Cxcl13
Cxcl12
G2M score
S score
num clones IGH
IGH MutFreq
num clones IGK
IGK MutFreq
Distance from LZ-DZ boundary (μm)
e
Bcl2a1b
G2M Score
f
TRB Clone
IGHG Clone
IGH CDR3 Length
IGHM Clone
g
Bcl2a1b
G2M Score
h
TRB Clone
IGHG Clone
IGH CDR3 Length
IGHM Clone
i
IGH-Secreted
Xbp1
Bst2
Selplg
Irf4
Mzb1
Edem1
H13
Cd40
Man1b1
Ddost
H2-Q7
j
IGH-Secreted
Xbp1
Bst2
Selplg
Irf4
Mzb1
Edem1
H13
Cd40
Man1b1
Ddost
H2-Q7
D14PI GC1
D21PI GC4

50

**Supplementary Figure 6. a.** Spatial maps show the segregated GCs from GraphST with the GC index on later timepoints LNs (D10PI, D14PI, D21PI). **b.** Segmentation of the D14PI GC1 into the light zone (blue: LZ) and dark zone (red: DZ). **c.** Boxplots show comparisons of marker genes (*Cxcl13*, *Cxcl12*, *Bcl2a1b*, *Aicda*, *Mfge8*) and G2M score between LZ (blue) and DZ (red) on the D14PI GC1. **d.** Line plots show LZ-DZ markers (*Cxcl13* and *Cxcl12*) LR features (number of clones and Mutation frequency of IGH and IGK) as a function of distance away from the LZ-DZ boundary; LZ direction (blue), DZ direction (red). **e&f.** Spatial map on D14PI GC1 on expression of LZ marker gene (*Bcl2a1b*) and G2M cell cycle scores, and adaptive immune repertoire information (TRB clone, length of IGH CDR3 in amino acid, number of IGHG/M clones) from LR. **g&h.** Spatial map on D21PI GC3 on expression of LZ marker gene (*Bcl2a1b*) and G2M cell cycle scores, and adaptive immune repertoire information (TRB clone, length of IGH CDR3 in amino acid, number of IGHG/M clones) from LR. **i&j.** Spatial map on genes positively correlated with *IGH-Secreted* (*Xbp1, Bst2, Selplg, Mzb1, Edem1, H13, Man1b1, Ddost, H2-Q7*), *Irf4*, and *Cd40* in Gaussian smoothing value.

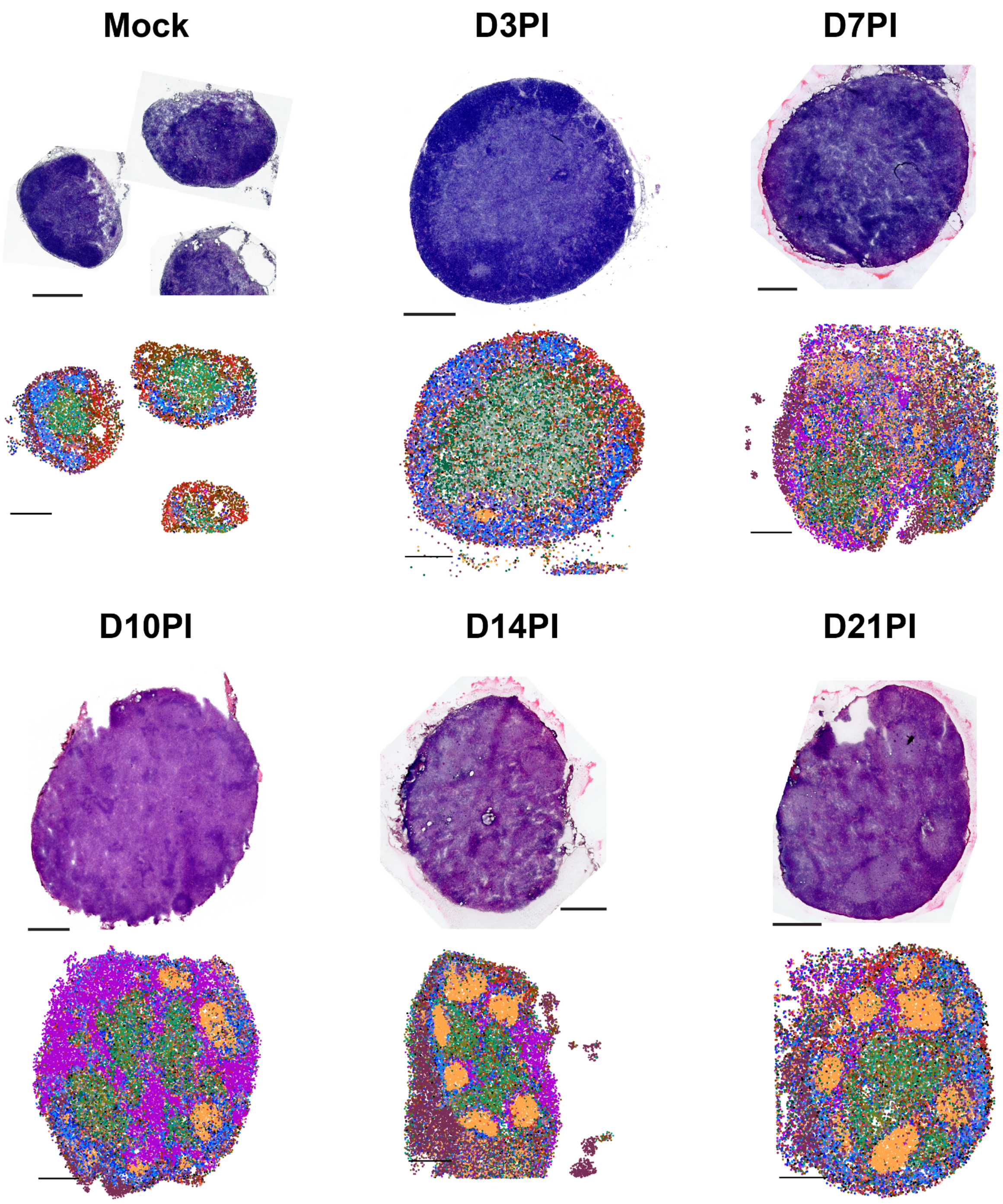


**Supplementary Figure 7.** Top row: hematoxylin and eosin (H&E) stained images of sister tissue sections from all the collected LN tissues (Mock, D3PI, D7PI, D10PI, D14PI, and D21PI). Bottom row: Spatial mapping of cell types across all sections, same as Figure 1.b.

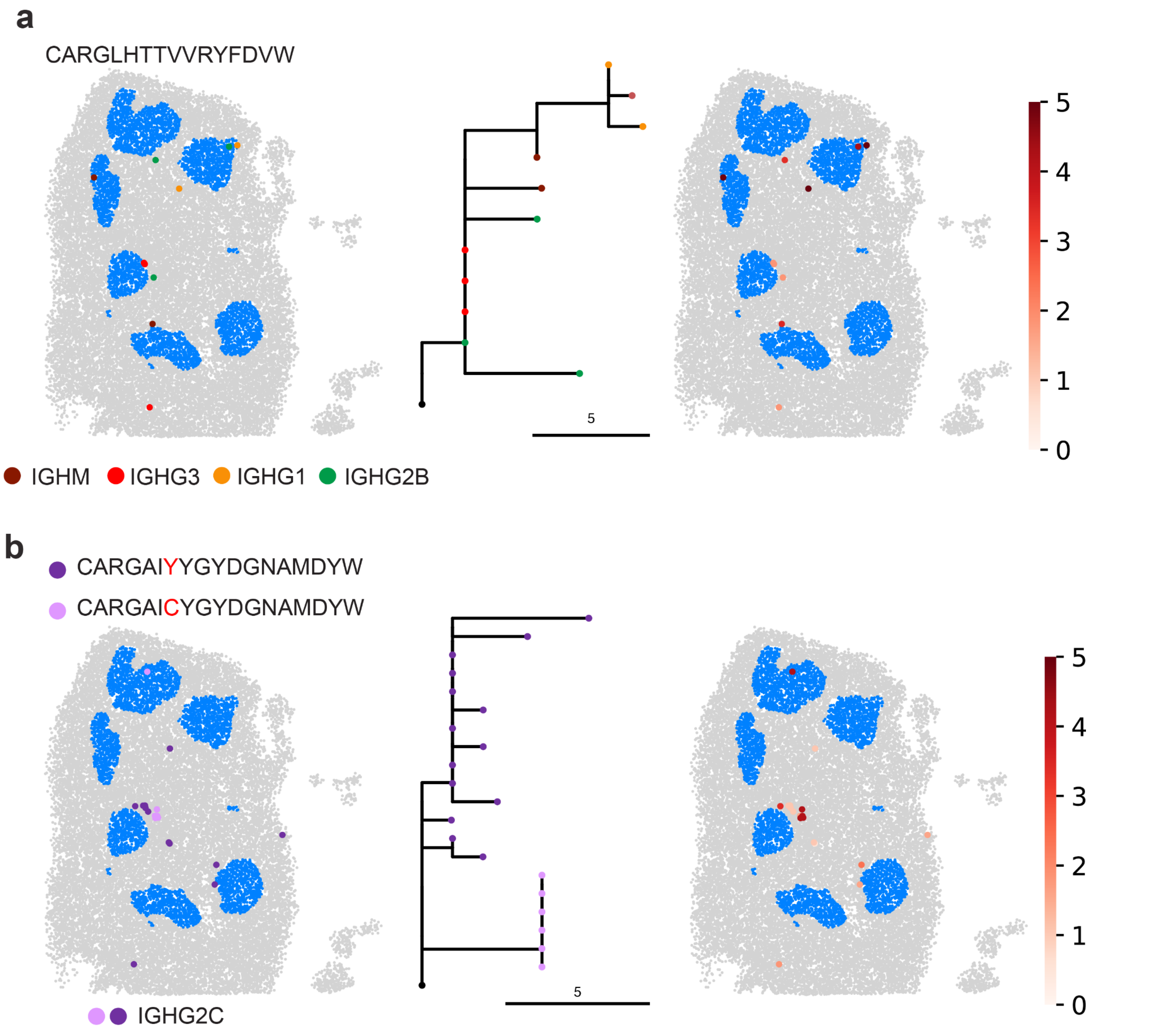


**Supplementary Figure 8.** Clonal evolution of two different IGH clone families from the D14PI sample. **a.** CDR3: CARGLHTTVVRYFDVW; **b.** CDR3: CARGAIYYGYDGNAMDYW. Spatial mapping and lineage tree represent each individual bead within the family, colored by different isoforms and CDR3 sequences. Branch lengths reflect the number of mutations (nt) accumulated from the germline (black node: root), and its spatial map.
